# Hunger and sleep recruit distinct brain systems to form spatial memory

**DOI:** 10.64898/2026.09.15.751731

**Authors:** Enea Tosadori, Joaquin Bou, Stoyan Dimitrov, Adrian Ocampo-Garces, Jose L. Valdes, Marion Inostroza, Jan Born, Anuck Sawangjit

## Abstract

By demonstrating that hunger consolidates spatial memory via a non-hippocampal mechanism that critically involves the retrosplenial cortex, we identify a mode of memory formation that is distinct from well-known hippocampus-dependent memory formation during sleep. Food-deprived rats encoded object locations and, during a subsequent 2-h consolidation phase, either received food and slept (Sleep-Fed) or stayed awake (Wake-Fed), or remained hungry and stayed awake (Wake-Hungry). At later retrieval testing, both Wake-Hungry and Sleep-Fed rats exhibited robust spatial memory, whereas Wake-Fed rats did not. Blocking neuropeptide-Y (NPY) signaling during the consolidation phase abolished hunger-consolidated memory. Unlike sleep-dependent consolidation, hunger-consolidated memory did not require the hippocampus during consolidation or retrieval, and not even during encoding. In contrast, inhibiting retrosplenial cortex during retrieval abolished hunger-consolidated memory but spared memory formed during sleep. Thus, recruitment of brain systems to form spatial memory is tuned to specific brain states, and fundamentally differs between sleep and hunger.

## Main

Memories, after encoding, undergo a period of consolidation to be maintained in the long-term^1^. The formation of long-term memory is commonly conceptualized as a systems consolidation process in which representations of new experiences are initially encoded into hippocampal networks that bind neocortical neuron assemblies representing the experience into a coherent memory trace^2,3^. Repeated reactivations of the initially encoded trace and overlapping traces transforms the initial representation such that neocortical components of the memory representation are strengthened and become independent of the hippocampus. Thus, spatial experience is thought to be obligatorily encoded initially into hippocampal networks. However, repeated reactivation of such hippocampal spatial traces strengthens neocortical components of such memories, for example, in retrosplenial cortex that then may also contain allocentric information necessary for navigation^2,4^.

Sleep is well known to support, and may even be critical for the systems consolidation of new information into long-term memory^5,6^. Since initial experiments by Rosa Heine^7^ numerous studies in humans and rodent models have shown that sleep after encoding benefits the later recall of different types of memories, in comparison with a post-encoding wake consolidation phase. Importantly, systems consolidation during sleep critically depends on hippocampal function, specifically on the repeated reactivation of newly encoded hippocampal traces preferentially occurring during slow wave sleep epochs^8–11^. In rats, suppression of hippocampal activity during a sleep consolidation phase immediately following encoding, abolishes memory for objects at a retrieval test taking place more than one week later, although the retrieval of these object memories does not require hippocampal function^8,12^.

While this evidence fostered the assumption that sleep could be the only brain state supporting the formation of long-term memory, initial findings in fruit flies suggest that memories may also be consolidated in the wake state, specifically when the animal is hungry^13^. In those experiments, starved flies formed persistent appetitive odor-food associations despite sleep loss. The hunger-driven consolidation process differed from sleep-associated consolidation in relying on neural pathways involving neuropeptide F, a homologue of the mammalian hunger signal neuropeptide Y (NPY)^14^. However, it remained unresolved whether hunger-dependent consolidation is a general process extending to non-appetitive memory, not specifically related to food stimuli, and whether it shares the common hippocampal mechanisms of sleep-dependent consolidation. Here, we compared in rats the consolidating effects of hunger and sleep in a spatial memory task, i.e., the classic object-place recognition (OPR) task which is thought to critically depend on hippocampal function^15,16^. We show that hunger consolidates OPR memory in awake rats and that, unlike sleep-dependent consolidation, hunger-dependent consolidation does not rely on hippocampal function, even for spatial memory. By contrast, retrieval of hunger-consolidated memory, but not sleep-consolidated memory, relies on the retrosplenial cortex.

## Results

### Hunger promotes consolidation of spatial memory during wakefulness

All animals were first food-deprived for 24 h to induce an intermediate level of hunger known to be associated with robust increases in brain NPY levels^17^ (see Figure 1a for experimental procedures). The period ended with the 10-min encoding phase of the object–place recognition (OPR) task. During the following 2-h experimental consolidation interval, the rats had either ad libitum access to food and were simultaneously allowed to sleep (Sleep-Fed group) or were kept awake (Wake-Fed group), or they remained food-deprived during the 2-h consolidation phase while awake (Wake-Hungry group). OPR retrieval was tested 24 h after encoding. Object exploration discrimination indices (DI) at the retrieval test indicated robust OPR memory in the Wake-Hungry and Sleep-Fed groups (Fig. 1b). DIs in both groups were significantly elevated already during the first 3 min of the retrieval phase, i.e., the period most sensitive to OPR memory effects^18–20^, and remained at this level throughout the entire 5-min phase (*p* < 0.002, Extended Data Fig. 1a). By contrast, the Wake-Fed group failed to express any OPR memory at retrieval testing (*p* > 0.177, Fig. 1b). Direct comparisons revealed that both the Sleep-Fed and Wake-Hungry groups performed better than the Wake-Fed group (*p* < 0.015, F(2,33) = 6.66, *p* = 0.004, for Group main effect, p = 0.293 for Group x Minute interaction), whereas performance was closely comparable between Wake-Hungry and Sleep-Fed groups (*p* = 1).

**Fig. 1.**
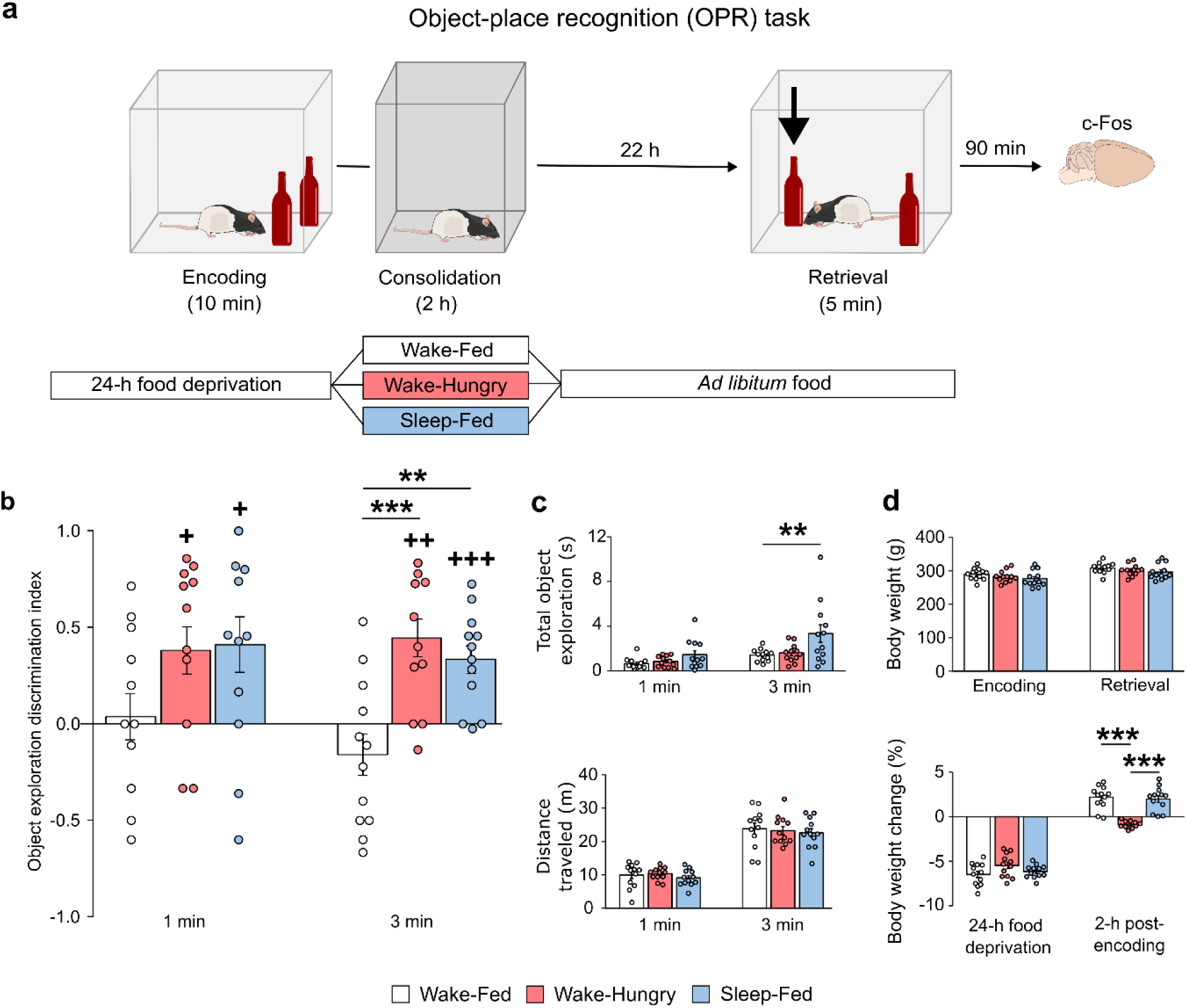
Hunger promotes consolidation of spatial memory during wakefulness. **a**, Experimental procedures: Rats were subjected to a 24-h food deprivation period which ended with the 10-min Encoding phase of the object-place recognition (OPR) task in which rats explored two identical objects in an arena surrounded by distal visual cues (not shown). Encoding was followed by the 2-h Consolidation phase in which the animals received food ad libitum and slept (Sleep-Fed group, blue) or remained awake (Wake-Fed group, white), or remained awake and continued to hunger (Wake-Hungry group, red). Following the consolidation phase, all groups had free access to food and were allowed to sleep. OPR memory was tested 22 h later during the 5-min Retrieval phase, in which rats explored the two familiar objects, with one of the objects being displaced from its original location (arrow). OPR memory is indicated by the object exploration discrimination index (DI), i.e., when the rat spends more time exploring the displaced than the non-displaced object. **b**, DIs at retrieval test. OPR memory is preserved in the Sleep-Fed and Wake-Hungry groups, but absent in the Wake-Fed group. **c**, Control measures of total object exploration time (*top*), distance traveled (*bottom*) at retrieval and **d**, body weight at encoding and retrieval phases (*top*) as well as changes in body weight after 24-h food deprivation and during the 2-h consolidation phase (*bottom*). Note, expected weight gain in Wake-Fed and Sleep-Fed groups, but not in Wake-Hungry group. N = 12 rats per group, data shown as mean + s.e.m. with overlaid dot plots. \*\*\**p* < 0.001, \*\**p* < 0.05, \**p* < 0.05 for pairwise two-sided *t*-test. ^+++^*p* < 0.001, ^++^*p* < 0.01, ^+^*p* < 0.05, for one-sample *t*-tests against chance level (see Extended Data Fig. 1a and 2a for analyses of the entire 5-min retrieval phase).

Control measures of total object exploration time during retrieval testing differed across the three groups (F(2,33) = 4.02, *p* = 0.028 for Group main effect), with the Sleep-Fed group showing a slight increase in total exploration time at 3 min of the test (F(2,33) = 5.17, *p* = 0.011, for Group x Minute interaction) which reached significance in comparison with the Wake-Fed group (*p* = 0.04, Fig. 1c *top*, Extended Data Fig. 2a). However, Sleep-Fed and Wake-Hungry groups did not differ in total object exploration (*p* > 0.057) and, importantly, including total object exploration time at 3 min as a covariate in a control analysis of DIs did not change the pattern of significant group differences observed for OPR retrieval performance (*p* = 0.008, for pairwise comparison between Wake-Hungry and Wake-Fed groups). The groups, moreover, did not differ in the distance traveled in the arena during retrieval (F(2,33) = 0.3, *p* = 0.747, for Group main effect, Fig. 1c *bottom*), ruling out that DI memory performance was driven by non-specific differences in locomotor activity or general arousal between groups.

Importantly, the rats of the Wake-Hungry group also did not differ in body weight from the two other groups at encoding or retrieval testing (all *p* > 0.29, Fig. 1d *top*). Analyses of changes in body weight, as an estimate of the animal’s hunger state, showed that the food provided during the 2-h post-encoding consolidation phase produced a significant gain in weight in the Sleep-Fed and Wake-Fed animals, in comparison with the Wake-Hungry animals which did not change in body weight (F(2,33) = 28.56, *p* < 0.001; *p* < 0.001, for pairwise comparisons with Wake-Hungry, Fig.1d *bottom*). Altogether, the findings in the Sleep-Fed and Wake-Fed conditions replicate numerous previous studies indicating that sleep after encoding is essential for forming lasting spatial memories on the OPR task^8,20–23^. Against this backdrop the retrieval performance of the Wake-Hungry group provides the first evidence that lasting OPR memory can also be formed in awake conditions when the animal is hungry.

### Neuropeptide Y mediates memory consolidation during hunger

In starved *Drosophila*, consolidation of appetitive odor-food association memory has been found to require neuropeptide F which is a homologue of the hunger signal neuropeptide Y (NPY) in the mammalian brain^13^. Against this backdrop, we asked whether consolidation of OPR memory in the Wake-Hungry condition likewise depends on NPY signaling. Two groups of rats underwent the same experimental procedures as the Wake-Hungry group of the main experiment except that they were intracerebroventricularly injected with either the NPY receptor antagonist BIBO 3304 (NPY-Y1R-Ant group) or saline solution (Saline group) prior to the 2-h post-encoding consolidation period. BIBO 3304 is a selective antagonist of the NPY-Y1 receptor (NPY-Y1R) with predominant post-synaptic expression^24^. Suppressing NPY signaling during the consolidation phase - during which the rats were awake and hungry - abolished the expression of OPR memory throughout the 5-min retrieval phase, whereas the Saline control group retained robust OPR memory (*p* > 0.255 and *p* < 0.037, for NPY-Y1R-Ant and Saline groups respectively, Extended Data Fig. 1b), with the two groups significantly differing in their memory performance (F(1,16) = 8.96, p = 0.009, for Group main effect, *p* = 0.249, for Group x Minute interaction; Fig. 2a). Control measures of total object exploration as well as distance traveled during the retrieval phase were comparable between the groups (*p* > 0.411, Fig. 2b, Extended Data Fig. 2b).

**Fig. 2.**
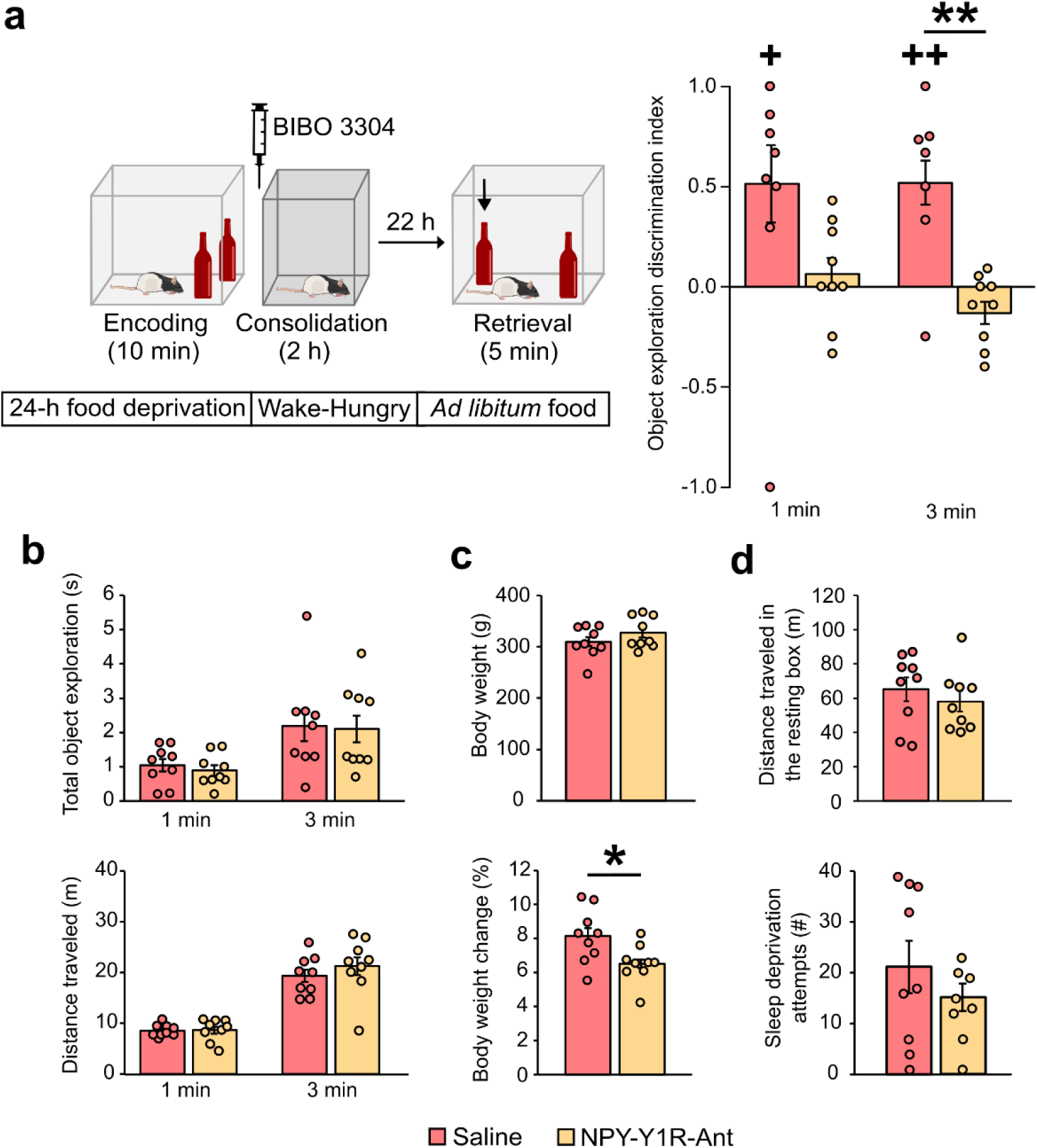
Neuropeptide Y mediates memory consolidation during hunger. *a, Left*, behavioral procedures were identical to that of the Wake-Hungry group of the main experiment (Fig. 1a), except that immediately after the OPR encoding phase, NPY-Y1 receptor antagonist (BIBO 3304) or saline was infused into the lateral ventricle. OPR memory was tested 24 h after encoding. *Right*, object exploration discrimination index (DI) in rats that received the NPY-Y1 receptor antagonist (NPY-Y1R-Ant, light gold bars) or Saline infusion (red bars). Note, absent OPR memory in NPY-Y1R-Ant group. **b**, Total object exploration (*top*) and distance traveled (*bottom*). **c**, Body weight at retrieval (*top*) and body weight change during the 22-h post-consolidation interval (*bottom*). Note, while in the NPY-Y1R-Ant group the increase in body-weight during the post-consolidation interval is reduced (consistent with the expected decrease in food intake following NPY-Y1R antagonist), absolute body weight at retrieval phase does not differ between groups. **d,** Distance traveled in the resting box (*top*) and number of sleep deprivation attempts (*bottom*) during the 2-h consolidation period were comparable between the groups. N = 9 rats per group. Data shown as mean + s.e.m. with overlaid dot plots, \*\**p* < 0.01, \**p* < 0.05 for pairwise two-sided *t*-test; ^++^*p* < 0.01, ^+^p < 0.05 for one-sample *t*-tests against chance level.

The NPY-Y1R-Ant rats, unlike the Saline controls, showed a significantly smaller increase in body weight from the end of the 2-h consolidation phase to the retrieval test (t(8) = −2.5, *p* = 0.024, Fig. 2c), likely reflecting reduced food consumption, consistent with the role of NPY in driving hunger^26^. Importantly, body weight at the OPR retrieval test did not differ between the NPY-Y1R-Ant rats and the Saline controls (*p* = 0.24, Fig. 2c). Moreover, there were no differences between the groups in locomotor activity or the number of gentle-handling interventions (to keep the animals awake) during the 2-h consolidation phase (*p* > 0.318; Fig. 2d), overall ruling out that the disruptive effect of blocking NPY signaling on OPR memory consolidation reflected substantial non-specific side effects of BIBO 3304 on general levels of arousal and stress^26^. Taken together, and aligning with findings in *Drosophila*^13^, these results support a critical role of NPY signaling in mediating memory consolidation in hungry conditions.

### Brain-state-dependent c-Fos expression during spatial memory retrieval

Considering that OPR retrieval performance was closely comparable between the Wake-Hungry and Sleep-Fed groups, we asked to what extent the animals in both groups recruited the same neuronal circuitry for successful retrieval of OPR memory. As an unspecific marker of neuronal activity, we analyzed expression of the immediate-early gene c-Fos during the retrieval phase^27^. We concentrated on cortical, thalamic, and hippocampal regions known from previous work to contribute to OPR memory^28^ and, additionally, on the hypothalamic arcuate nucleus and lateral hypothalamus as areas involved in mediating hunger signaling also in the context of learning processes^29^. In hippocampal and parahippocampal areas expression of c-Fos during retrieval was highest in the Sleep-Fed group, with this activation being restricted to selected areas like the perirhinal (PRh) cortex (*p* < 0.017, compared with the other two groups), entorhinal cortex (Ent), dentate gyrus (DG) and CA1 (*p* < 0.031, compared with home cage control levels; F(8,132) = 2.36, *p* = 0.02, for Group x Region interaction, Fig. 3a). In medial prefrontal cortical areas c-Fos expression was also highest in the Sleep-Fed group, in particular in the prelimbic region (PL; *p* = 0.012, compared with Wake-Hungry group) and cingulate gyrus (CG; *p* = 0.034, F(4,66) = 3.9, *p* = 0.007, for Group x Region interaction, Fig. 3b). Such recruitment of hippocampal and medial prefrontal cortex during successful OPR retrieval as observed here in the Sleep-Fed group. has been similarly observed in previous studies^28^. Notably, the Wake-Hungry group did not show any c-Fos levels above those of the home cage control condition in the two hypothalamic target areas, the arcuate nucleus and lateral hypothalamus, c-Fos levels were even significantly diminished compared with both the Sleep-Fed and Wake-Fed groups (F(2,33) = 9.73, *p* < 0.001, for Group main effect, *p* < 0.002, for post-hoc pairwise comparisons, Fig. 3c). No differences between groups were found in posterior cortical and thalamic regions (all *p* > 0.113, Extended Data Fig. 3).

**Fig. 3.**
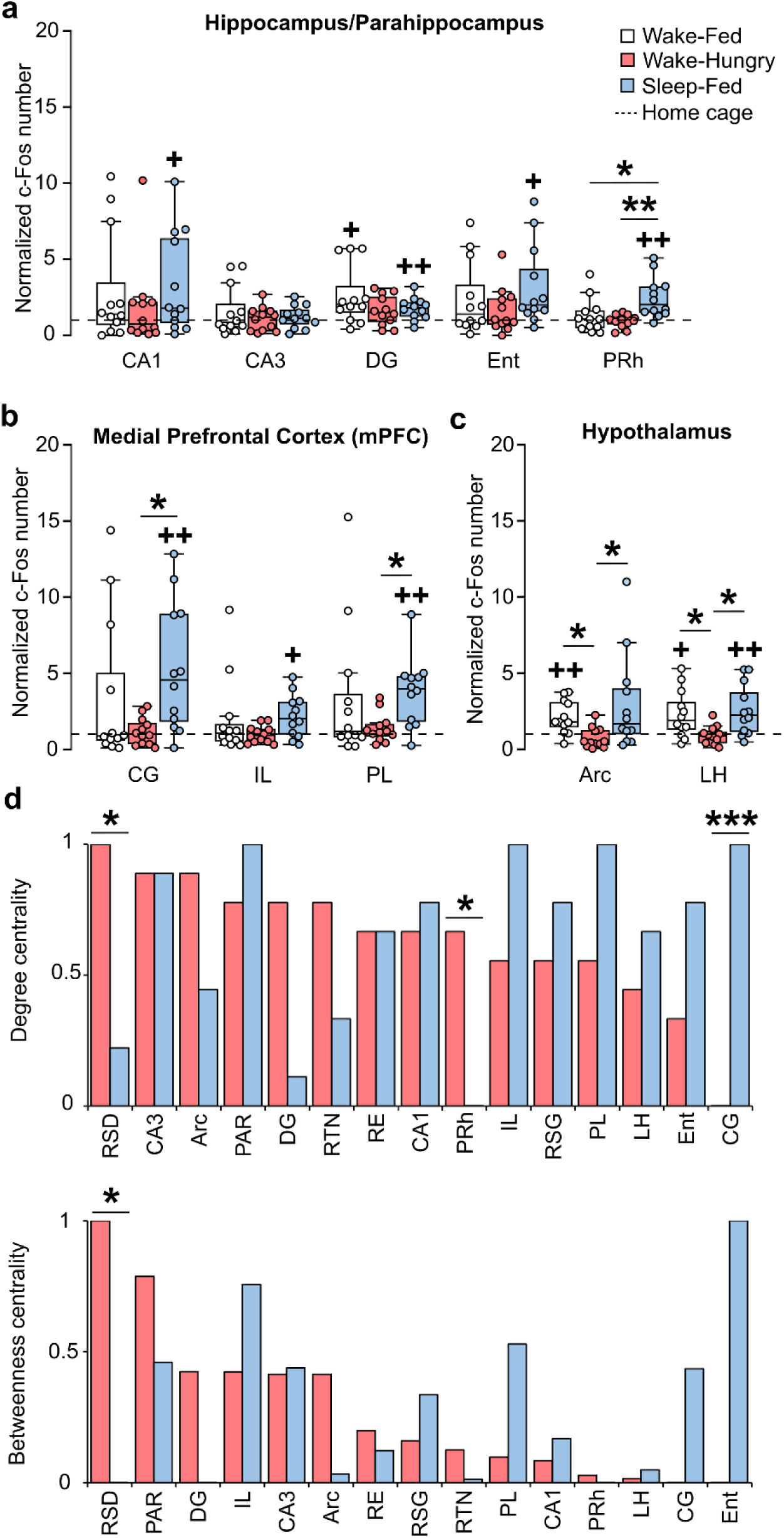
Brain-state dependent c-Fos expression during spatial memory retrieval. Number of c-Fos positive cells relative to home cage control group (set to 1, dashed lines) at OPR retrieval phase in (**a)** hippocampal/parahippocampal areas in the Wake-Fed (white), Wake-Hungry (red), and Sleep-Fed (blue) groups of the main experiment. In Sleep-Fed animals, c-Fos expression was higher than in both Wake-Hungry and Wake-Fed groups in PRh, and upregulated in CA1, DG and Ent compared to home cage controls, consistent with generally enhanced activation levels in hippocampal complex. **b**, c-Fos expression in medial prefrontal cortex (mPFC) regions. Note, higher c-Fos expression in Sleep-Fed than Wake-Hungry group in CG and PL. **c**, c-Fos expression in hypothalamus. Note, reduced c-Fos expression in Wake-Hungry group in both Arc and LH compared with Sleep-Fed and Wake-Fed groups. **d**, Functional network analysis based on interregional correlations of c-Fos expression during OPR retrieval (Extended Data Fig. 3): normalized degree centrality (*top*) and betweenness centrality (*bottom*) across brain regions for Sleep-Fed (*blue*) and Wake-Hungry (*red*) groups. Note, opposing patterns of both centrality metrics between groups: Sleep-Fed group shows higher centrality in medial prefrontal areas (CG, PL, IL) and Ent, whereas RSD and PAR were the most prominent hub areas in Wake-Hungry group. Values were normalized to the minimum and maximum within each group with ordering of regions (from highest to lowest) according to the Wake-Hungry group. N = 12 rats per group. Data shown in a-c as boxplots with overlaid dot plots. Boxes represent the median and interquartile range (IQR), whiskers extend to 1.5 x IQR. \**p* < 0.05, \*\**p* < 0.01, *** *p* < 0.001, for pairwise comparisons between groups (in a-c two-sided *t*-test, in d permutation-based Network Comparison Test). ^++^*p* < 0.01, ^+^*p* < 0.05 for a one-sample Wilcoxon signed-rank test against home cage control. Abbreviations: Arc = arcuate nucleus, CA1 and CA3 = cornu ammonis, CG = cingulate gyrus, DG = dentate gyrus, Ent = entorhinal cortex, IL = infralimbic cortex, LH = lateral hypothalamus, PAR = parietal cortex, PRh = perirhinal cortex, PL = prelimbic cortex, RE = nucleus reuniens, RSD = retrosplenial dysgranular cortex, RSG = retrosplenial granular cortex, RTN = reticular nucleus of the thalamus.

To further explore the differential brain activation profiles for retrieving OPR memory in the Wake-Hungry and Sleep-Fed groups, we performed a functional network analysis based on interregional correlations of c-Fos expression during the retrieval phase^30^. Networks were summarized with two centrality metrics: *degree centrality* indexing how broadly a region is coactive with others (Fig. 3d, *top*), and *betweenness centrality* indexing how often a region lies on shortest paths between regions and may thus act as a communication bridge (Fig. 3d, *bottom*). While overall degree centrality (*p* = 0.754) and betweenness centrality (*p* = 0.939) of the functional networks were comparable between Wake-Hungry and Sleep-Fed groups, functional networks differed in quality: In the Sleep-Fed group, medial prefrontal cortical areas (CG, PL, IL) together with the entorhinal cortex (Ent) were most highly connected nodes during OPR memory retrieval, whereas these regions showed no comparable functional connectivity in the Wake-Hungry group, with a highly significant difference in degree centrality between groups for CG (Fig. 3d, *top*, C = –10, p < 0.001). In sharp contrast, for the Wake-Hungry group the dysgranular retrosplenial cortex (RSD) emerged as a dominant hub area during OPR retrieval showing both distinctly higher degree and betweenness centrality than in the Sleep-Fed group (C = 6, *p* = 0.03 and C = 15.1, *p* = 0.037 respectively, Fig. 3d). Additionally, degree centrality of the perirhinal cortex (PRh) was enhanced in this group (C = 5, *p* = 0.043, Fig. 3d, *top*). Overall, the patterns suggest that despite their highly comparable behavioral performance, Sleep-Fed and Wake-Hungry groups engage different brain networks for OPR memory retrieval, relying on medial prefrontal and retrosplenial hub areas, respectively.

### Hippocampus is not critical for spatial memory consolidated during hunger

Consolidation of memory during sleep is known to critically rely on hippocampal function, specifically on the neuronal replay of newly encoded hippocampal representations^10,31^. The hippocampal dependency of consolidation during sleep pertains both to memories that are ultimately stored outside the hippocampus and can later be retrieved independently of it as well as to spatial memory traditionally considered to rely on hippocampal networks^8–11^. We therefore asked whether OPR consolidation during hunger would likewise rely on hippocampal function. The question was particularly pressing given that our c-Fos analyses pointed to a disengagement of the hippocampus during OPR retrieval when these memories had been consolidated during hunger (Fig. 3). In two separate experiments we compared the effects of inhibiting hippocampal function during the 2-h post-encoding consolidation phase and during the retrieval phase, respectively, between OPR memory consolidated during sleep and during hunger (Fig. 4a). In the first “consolidation” experiment, rats were tested twice separated by a 1-week interval, once exposed to the Wake-Hungry condition, and once to the Sleep-Fed condition. The experimental conditions were identical to those of the respective groups of the main experiment, except that hippocampal function was reversibly inactivated during the consolidation phase using a bilateral injection of the GABA_A_ receptor agonist muscimol, into the dorsal hippocampi. At the retrieval test 24 h later, rats did not show any OPR memory with the hippocampi inactivated during a consolidation phase spent in the Sleep-Fed condition (*p* > 0.583). By contrast, when the hippocampus was inactivated during a Wake-Hungry consolidation phase, the rats showed a well preserved OPR memory at retrieval testing 24 h later (*p* < 0.001, F(1,8) = 23.11, *p* = 0.001, for the difference between Sleep-Fed vs Wake-Hungry conditions, Fig. 4b, *left*, Extended Data Fig. 1c, *left*). Control measures of total exploration time and distance traveled during retrieval were comparable between the conditions (*p* > 0.132 and *p* > 0.255, Fig. 4c, Extended Data Fig. 2c, *left*). The findings indicate that unlike sleep-dependent consolidation, consolidation of OPR memory during hunger in awake rats does not require hippocampal function.

**Fig. 4.**
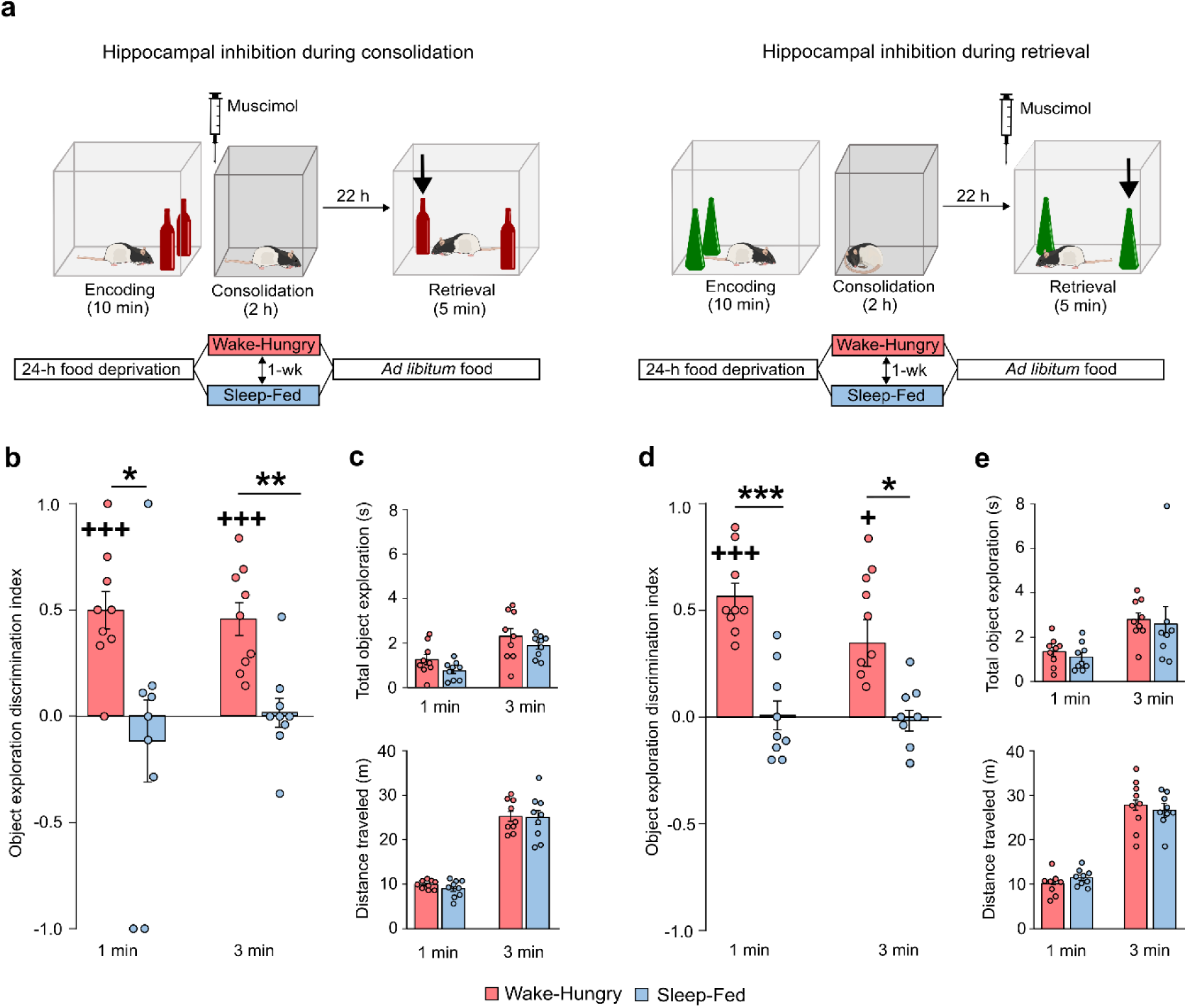
Hippocampus is not critical for forming and retrieving spatial memory consolidated in the Wake-Hungry condition. **a,** Behavioral procedures were identical to those of the main experiment (see Fig. 1a), except that in one experiment, hippocampus was inhibited during the 2-h Consolidation phase (*left*) whereas in the other experiment it was inhibited during the Retrieval phase 24 h later (*right*). Reversible inhibition was achieved by bilateral injection of the GABA_A_ receptor agonist muscimol into the dorsal hippocampus (syringe) before the respective phase. Both experiments were designed as within-subject comparisons with each rat tested once on the Wake-Hungry condition (*red bars*) and another time on the Sleep-Fed condition (*blue bars*), separated by 1 week (n = 9 rats per experiment). **b** & **d**, Object exploration discrimination index (DI) at retrieval test in the Wake-Hungry and Sleep-Fed conditions. Note, whereas hippocampal inhibition during consolidation (b) and retrieval (d) abolished OPR memory when consolidated in Sleep-Fed conditions, it was well-preserved when consolidated in Wake-Hungry conditions. **c** & **e,** Control measures of total object exploration time (*top*) and distance traveled (*bottom*) during the retrieval phase did not differ between Wake-Hungry and Sleep-Fed conditions in both experiments. Data shown as mean + s.e.m. with overlaid dot plots, ***p < 0.001, \*\**p* < 0.01, \**p* < 0.05 for pairwise two-sided *t*-test. ^+++^*p* < 0.001, ^+^*p* < 0.05 for one-sample *t*-test against chance level.

Procedures in the second “retrieval” experiment were identical to those of the first, except that the dorsal hippocampi were inactivated during the retrieval phase. Again, when OPR memory had been consolidated during a Sleep-Fed consolidation phase, blocking the hippocampus during retrieval abolished OPR memory expression (*p* > 0.758, Fig. 4d), replicating previous studies^15,16,32^. By contrast, when the OPR memory had been consolidated in the Wake-Hungry condition, the rats showed a well preserved OPR memory at the retrieval test 24 h later (*p* < 0.017, F(1,8) = 38.01, *p* < 0.001, for the difference between Sleep-Fed vs Wake-Hungry conditions, Fig. 4d). Total object exploration time and distance traveled during the retrieval phase did not differ between the conditions (*p* > 0.429 and *p* > 0.233, respectively, Fig. 4e). These results indicate that, unlike sleep, hunger consolidates a spatial memory representation that likely resides outside the hippocampus.

Given the hippocampal independence of OPR memory consolidated during hunger, in a third experiment, we examined whether formation of these spatial memories in the hungry state relies on representations that were already *encoded* outside the hippocampus. Experimental procedures, here, remained the same, except that the dorsal hippocampi were inactivated during the OPR encoding phase and that we used optogenetic inhibition (via bilateral pAAV-hSyn-JAWS-KGC-GFP-ER2 virus injection, light-ON 8s, light-OFF 2–2.5s throughout encoding), in order to precisely restrict inactivation to the 10-min encoding phase leaving undisturbed the subsequent consolidation phase (Fig. 5a). Corresponding to the foregoing experiments, inactivating the hippocampus during encoding abolished any OPR memory expression at the retrieval test 24 h later, when memories were consolidated in the Sleep-Fed condition (*p* > 0.485, Fig. 5b), indicating that rats relied on a hippocampally encoded spatial representation for retrieval performance. By contrast, rats consolidating memory in the Wake-Hungry condition, despite hippocampal inactivation at encoding, still expressed a full-blown OPR memory at the retrieval test (*p* < 0.025, F(1,6) = 6.7, *p* = 0.041, for the difference between Sleep-Fed vs Wake-Hungry conditions, F(1,6) = 0.09, *p* = 0.771, for Condition x Minute interaction, Fig. 5b). Total object exploration and distance traveled at retrieval testing did no differ between conditions (all *p* > 0.761 and *p* > 0.154, respectively, Fig. 5c). These findings are consistent with the view that rats encode multiple - hippocampal and extrahippocampal - representations with post-encoding sleep consolidating the hippocampal representations, whereas hunger selectively consolidates the extrahippocampal representations.

**Fig. 5.**
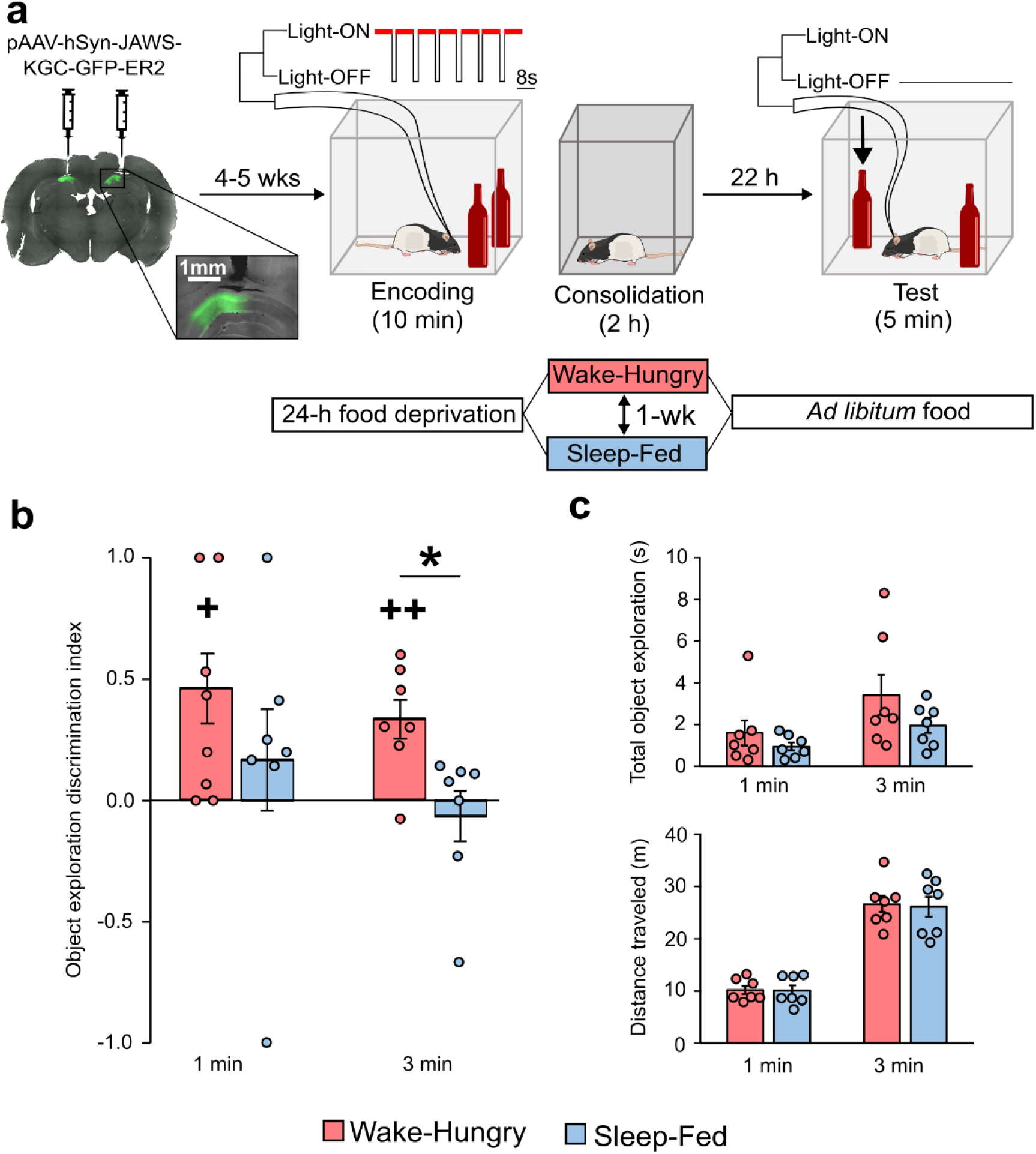
Hippocampus is not critical for encoding spatial memory consolidated in the Wake-Hungry condition. **a,** Behavioral procedures and experimental design were identical to those in Fig. 4a except that the hippocampus was inhibited during the 10-min Encoding phase. Reversible inhibition was achieved optogenetically: pAAV-hSyn-JAWS-KGC-GFP-ER2 virus was injected bilaterally into dorsal hippocampal CA1 (*green)* 4-5 weeks before experiments. During the OPR encoding phase, 625nm LED light (15 mW) was delivered (light-ON 8s, light-OFF 2–2.5s). **b**, Object exploration discrimination index (DI) at retrieval test in the Wake-Hungry and Sleep-Fed conditions. Note, whereas hippocampal inhibition during encoding abolished OPR memory when consolidated in Sleep-Fed conditions (*blue bars*), it was well-preserved when consolidated in Wake-Hungry conditions (*red bars*), despite hippocampal suppression during the encoding phase. **c**, Total object exploration time (*top*) and distance traveled (*bottom*) during the retrieval phase did not differ between conditions. N = 7 rats, data shown as mean + s.e.m. with overlaid dot plots, \**p* < 0.05 for pairwise two-sided *t*-test. ^++^*p* < 0.01, ^+^*p* < 0.05 for one-sample *t*-test against chance level.

### Retrosplenial cortex is required for retrieving spatial memory consolidated during hunger

Our functional network analyses pointed to the dysgranular retrosplenial cortex (RSD) as a potential hub area regulating retrieval of OPR memory after hunger-dependent consolidation whereas this area appeared to be disengaged at retrieval after sleep-dependent consolidation (Fig. 3d). To test a causal contribution of the RSD to OPR memory, in a further experiment we bilaterally inactivated the RSD during the OPR retrieval phase via muscimol injection using the same procedures as described for the corresponding experiments with suppression of hippocampal activity (Fig. 6a, Extended Data Fig. 4c). Blocking RSD during the retrieval phase completely abolished the expression of OPR memory when the memory had been consolidated during a Wake-Hungry consolidation phase (*p* > 0.404), but left OPR memory intact when the memory had been consolidated during sleep (*p* < 0.013, F(1,5) = 14.25, *p* < 0.013, for difference between Sleep-Fed and Wake-Hungry conditions, Fig. 6b). Control measures of total object exploration time during retrieval testing did not differ between conditions (*p* > 0.59, Fig. 6c, *top*). Rats in the Sleep-Fed condition traveled a longer distance as compared to the Wake-Hungry condition during the first minute of the test phase (*p* = 0.009, F(1,5) = 0.58, *p* = 0.441, for Condition x Minute interaction), but this transient difference was not maintained throughout the test phase (*p* = 0.535, Fig. 6c, *bottom*, Extended Data Fig. 2d). Moreover, the difference in DI memory performance between groups was confirmed in a control analysis including distance traveled at 1 min as covariate (F(1, 17.48) = 13.18, *p* = 0.001), overall excluding that systematic locomotor differences during retrieval confounded memory performance. These results indicate that, unlike sleep, hunger consolidates OPR memory into representations that depend on the RSD, taking over a role in regulating spatial memory retrieval similar to that previously thought to critically rely on hippocampal function^33,34^.

**Fig. 6.**
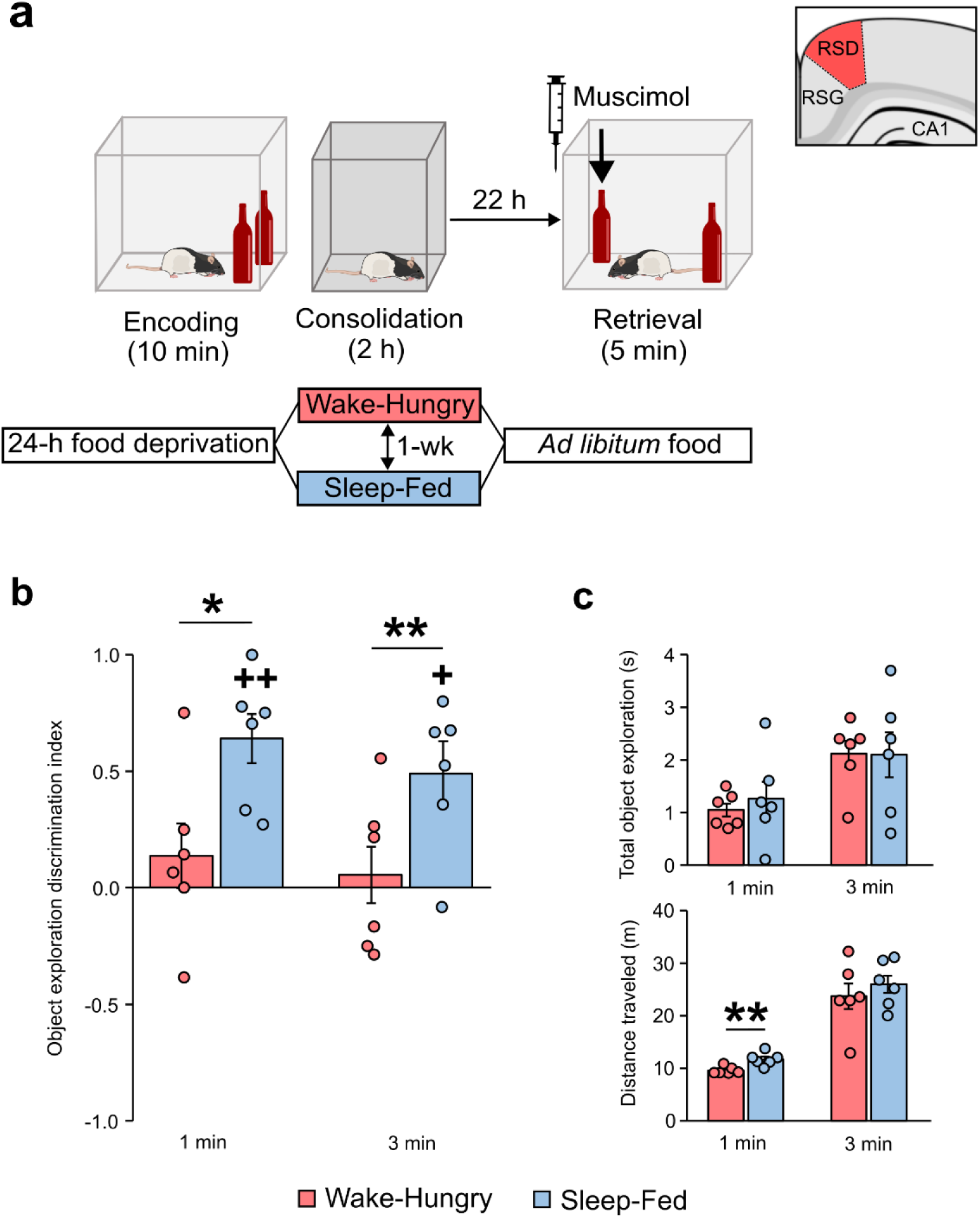
Retrosplenial cortex is critical for retrieving spatial memory consolidated in the wake-hungry condition, but not after sleep consolidation. **a,** Behavioral procedures and experimental design were identical to those in Fig. 4a except that the dysgranular retroplenial cortex was inhibited during the OPR retrieval phase. Reversible inhibition of the RSD was achieved pharmacolocially (bilateral injection of muscimol). **b**, Object exploration discrimination index (DI) at retrieval test in the Wake-Hungry (*red*) and Sleep-Fed (*blue*) conditions. **c**, Total object exploration time (*top*) and distance traveled (*bottom*) during the retrieval phase. The transient increase in locomotion in the Sleep-Fed animals at 1 min was unrelated to DI memory performance (Extended Data Fig. 2d). N = 6 rats. Data shown as mean + s.e.m. with overlaid dot plots \*\**p* < 0.01, \**p* < 0.05 for pairwise two-sided *t*-test. ^++^*p* < 0.01, ^+^*p* < 0.05 for one-sample *t*-tests against chance level.

## Discussion

Our experiments provide a classical double dissociation between memory systems^35^ indicating that spatial experience can be encoded, consolidated into long-term stores, and retrieved from memory using two different brain systems, a hippocampus-dependent system and a retrosplenial cortex-dependent system. Whereas the hippocampus-dependent system forms long-term memory during sleep, the retrosplenial cortex-dependent memory system forms long-term memory in the wake state when the individual is hungry. The findings support the view of multiple traces encoding an experience^2,36–39^, with the specific traces and systems forming persistent memory depending on the organism’s state, such as sleep and hunger.

Whereas previous research indicates that the formation of long-term spatial memory in rodents as well as in humans requires sleep after encoding^20,22,40,41^, here we show that spatial memory can also be consolidated during a post-encoding wake period, however, only when the animal is hungry. Having ad libitum access to food during the post-encoding wake period prevented spatial memory consolidation. This finding aligns with growing evidence suggesting facilitating effects of hunger on memory formation^42–46^. Thus, starved *Drosophila* showed enhanced retention of odor-food associations^13^. Extending those findings in *Drosophila* we demonstrate here that the strengthening effects of hunger also pertains to non-food-related spatial stimuli. We, moreover, show that the consolidating effect of hunger depends on NPY transmission, i.e., an evolutionary preserved and highly specific orexinergic signal that is synthesized in the arcuate nucleus of the hypothalamus and acts mainly via widespread Y1 and Y5 receptors to stimulate food intake^47^. Inhibiting NPY-Y1 receptors specifically during the post-encoding consolidation phase abolished OPR memory in the awake rats when they were hungry, consistent with findings in *Drosophila*^13^.

At the same time, the effects observed after blocking NPY-Y1 receptors can be taken to rule out that consolidation processes in our experiments were confounded by non-specific stress-related factors^48,49^. We used only moderate levels of food deprivation to induce hunger and gentle handling to keep animals awake for a relatively short 2-h period, i.e., procedures well known to minimize stress^50,51^. Importantly, NPY is known to robustly reduce stress and anxiety, with the inhibition of NPY transmission leading to an increase, rather than decrease in stress levels^52,53^. Therefore, if stress-related mechanisms were driving memory enhancements, memory performance should have been higher in rats receiving the NPY-Y1 receptor antagonist. However, the antagonist instead entirely suppressed memory consolidation.

Crucially, we demonstrate that hunger and sleep rely on distinct neuronal systems to form spatial long-term memory. Consolidation during sleep requires hippocampal networks and was linked to enhanced retrieval-related (c-Fos) activity in canonical prefrontal perirhinal, and hippocampal areas well-known to regulate spatial memory processing^54–56^. Indeed, our findings that inhibiting hippocampal function during the experimental sleep consolidation phase - as well as inhibitions induced during encoding and retrieval phases - corroborates the active systems consolidation concept of sleep-dependent memory formation that, based on multiple findings, proposes hippocampal replay to play a key role in consolidating memory regardless of their eventual dependence on hippocampal function at retrieval^6,8,9,11^. In contrast to sleep-dependent consolidation, hunger-mediated consolidation requires the dysgranular retrosplenial cortex while hippocampal areas remain disengaged and retrieval-related activity in the hypothalamic areas appears to be even suppressed.

Paradoxically, after hunger-mediated consolidation rats were able to retrieve the OPR memory regardless of whether hippocampal function had been inhibited during consolidation, and memory performance was also not altered in these animals when hippocampal function was inhibited during the retrieval phase, indicating that hunger forms a hippocampus-independent spatial representation that is used for retrieving object locations. This is a surprising finding against the backdrop that the hippocampus is considered to be crucial for spatial memory and the regulation of behavior based on allocentric spatial mapping as required on our OPR task^16,57,58^. To the best of our knowledge, the present study is the first to demonstrate in rodents that the brain can successfully form and retrieve spatial object-location memories in the absence of hippocampal function during consolidation and retrieval^15,59^. These spatial memories were preserved even when optogenetic hippocampal inhibition was precisely restricted to the encoding phase, indicating that hunger-mediated consolidation accesses extrahippocampal spatial traces. Given that identical optogenetic inhibition during encoding prevented subsequent sleep-dependent consolidation of the spatial memories, our data provide evidence that the brain encodes different traces of the same spatial experience that are used for consolidation during sleep and hunger, respectively^36,38^.

Interestingly, the dysgranular retrosplenial cortex (RSD) emerged as a hub mediating retrieval of spatial memories when these were consolidated during hunger. The RSD has dense reciprocal connections with various cortical and sub-cortical brain structures including the hippocampus^4,60^. Retrosplenial cortex and hippocampus are commonly thought of as parallel systems regulating spatial navigation either in a complementing or contrasting manner based on types of learning or input information^61,62^. Consistent with the present findings, lesions to the RSD impaired spatial navigation in several studies^63,64^ overall suggesting that retrosplenial cortical networks are capable of supporting some form of allocentric spatial mapping. Different from these foregoing studies, here we concentrated on the consolidation of long-term spatial representations, providing novel evidence that hippocampal spatial representations formed during sleep can compensate for deficits in long-term retrosplenial cortical representations.

Taken together, our experiments identify distinct memory systems linked to the formation of spatial long-term memory during sleep and hunger, respectively, which aligns with ideas of multiple traces in the brain representing a specific experience^2,36,39^. What is the adaptive function of such dual memory systems seemingly consolidating similar experience in parallel? And, do these systems compete or synergize^65^? These are emergent questions. Hunger mediating long-term memory points to an evolutionarily conserved mechanism that, bypassing hippocampus-dependent consolidation during sleep, may allow for more flexible memory formation fine-tuned to metabolic need. A sleep-independent pathway of memory formation may, for example, help store possibly very long routes taken in the search for food in the long term without having to switch to sleep^66^ and, in this vein, may likewise help compensate for memory deficits in clinical conditions, such as Alzheimeŕs disease, where hippocampus-dependent memory consolidation during sleep is impaired^67,68^.

## Methods

### Animals

102 adult male Long Evans rats (Janvier, Le Genest-Saint-Isle, France, 250-350g, 12-15 weeks at test phase) were used for the experiments. Rats were paired-housed and kept at constant room temperature (22 ± 1° C) under a 12-h light/12-h dark cycle (lights on at 7:00 am). Rats had unrestricted access to water and food except during the experimental sessions. All experimental procedures were performed in accordance with European animal protection laws and were approved by the Baden-Württemberg state authority.

### Design and general procedures

A total of 6 experiments were conducted: (1) The main experiment including the Wake-Fed, Sleep-Fed and Wake-Hungry groups (n =12 rats per group), (2) the NPY-Y1 receptor antagonist (NPY-Y1R-Ant) experiment including the NPY-Y1R-Ant and Saline control groups (n = 9 rats per group), three Hippocampal inactivation experiments where inactivation was pharmacologically induced during (3) the experimental Consolidation phase (n = 9 rats), (4) the Retrieval phase (n = 9 rats), or (optogenetically induced) during (5) the Encoding phase (n = 7 rats), and (6) a RSD inactivation experiment (n = 6 rats) where the dysgranular retrosplenial cortex was pharmacologically inhibited during the Retrieval phase of the OPR task. Whereas the main experiment and the NPY-Y1R antagonist experiment followed a group design, the remaining experiments (3-6) were designed as within-subject comparison with each rat subjected to two conditions equivalent to the Sleep-Fed and Wake-Hungry conditions of the main experiment, with the order of the two conditions balanced across animals. The two conditions for each animal were separated by an interval of 1 week: In experiments 1 and 2, rats were randomly assigned to experimental groups before the experiment. The experimenters were blinded to the experimental group or condition during all offline analyses of behavioral recordings and c-Fos analyses, but not during behavioral data collection. Behavioral experiments took place during the animal’s rest phase (between 8:00 am and 3:00 pm). The main experiment for all animals comprised a 24-h food deprivation period that ended with the encoding phase of the OPR task. Encoding was followed by a 2-h experimental Consolidation phase and a delayed Retrieval phase 22 h later (Fig. 1a). Experiments were performed in a room equipped with a constant masking-noise generator and maintained at stable temperature (22± 1 °C) to minimize external disturbances.

#### Handling, habituation and food deprivation

Prior to the experiments, rats were handled for 5 min per day, at least once daily for five consecutive days, to habituate them to the experimenter and to general procedures. Animals also underwent a three-day habituation to the experimental context. Each day, the rat was placed into an empty standard rat cage with an object (not used in the experiments) inside and allowed to explore it freely for 10 minutes. Afterward, the rat was placed in the empty open field (80x80 cm, height: 40 cm) surrounded by distal cues, facing a different wall at each session to facilitate allocentric navigation. The rat was allowed to explore the arena for 10 minutes. Following arena familiarization, the rat was left undisturbed in the resting box (35x35 cm, height: 45 cm) containing some bedding material for 2-4 h.

After the last habituation session, animals were transferred to a clean home cage containing no food. The cage was replaced to prevent that small residual food fragments or odor cues could interfere with the intended food-deprivation manipulation. Water was available *ad libitum* throughout the entire deprivation period. To monitor body weight and ensure compliance with ethical standards, animals were weighed at four predefined time points: immediately after the last habituation day, prior to the encoding phase, after the 2-h experimental consolidation period and after the retrieval test.

#### OPR memory test

The OPR task comprised a 10-min encoding phase and a 5-min retrieval phase, 24 h later. During encoding, rats explored two identical objects. The positions of the objects were randomized across rats to avoid position bias. For the retrieval test, one of the two objects from the encoding phase was moved to a different location in the arena and each rat was given 5 minutes to explore the arena and objects. To enforce allocentric spatial mapping, rats were placed in the arena facing different walls during the encoding and retrieval phases. Immediately after encoding, animals were placed in the resting box for the 2-h consolidation phase. In conditions where food was available right after encoding (i.e. Sleep-Fed and Wake-Fed), the standard rat chow (22-31 g) were placed inside the resting box. In the sleep condition, during the 2-h consolidation phase, the animals were left undisturbed in the resting box, containing some bedding materials. In the wake conditions, wakefulness was enforced using gentle handling^12^. This procedure minimizes stress and confounding influences of locomotion. It consisted of tapping on the box and, if necessary, gently shaking the box. Video records ensured that signs of startle or freezing did not occur. Water was available *ad libitum* within the resting box for all groups. At the end of the 2-h consolidation phase, all animals were transferred to their home cages, where they had ad libitum access to food and water.

Objects used in the OPR task were made of glass with different colours and shapes and were sufficiently heavy to prevent displacement by the animals (height: 16-30 cm, base diameter: 7-11 cm). The two identical objects were positioned near the corner, each placed 15 cm from the adjacent walls and equidistant from the corner to prevent the animal’s preference for corner areas from biasing exploration. Each rat’s exploration behavior was recorded using a video camera and analyzed offline by an experienced researcher with ANY-maze software (Stoelting, Europe). After each experimental phase, the arena and objects were cleaned with water containing 70% ethanol to prevent odor cues from influencing subsequent sessions.

#### Procedures for remaining experiments

For the NPY-Y1R antagonist experiment (Experiment 2) and the Hippocampal and the RSD inactivation experiments (Experiment 3−6), procedures were identical to those described for the main experiment (Experiment 1), except that during habituation sessions animals were also habituated to the substance infusion procedure (Experiment 3, 4 and 6) or the optic cable-connection procedure (Experiment 5), in addition to habituation to the object, arena and resting box. Habituation to the arena and to the drug infusion / optogenetic manipulation procedures followed the same temporal order as in the actual experiments. For the optogenetic experiment (Experiment 5), animals during habituation were connected to the optic-fiber patch cables but the light source remained switched off. The cables were attached to a compensatory counter-weight system to facilitate the animal’s flexible navigation in the arena and minimize behavioral interference. Animals were connected to the cables during both the encoding and retrieval phases to maintain the same tethered condition across sessions.

In experiments 3−6 designed as within-subject comparisons, the second condition was always conducted in a context different from that of the first condition (i.e. different arena, distal cues, and resting box) to minimize potential interference from the contextual information with repeated testing. Habituation in the second context followed the same procedure as in the first, except that object habituation was not included, and a different set of objects was used for the OPR task.

### Analysis of memory performance

Exploration was defined by the rat being within 1 cm of an object, directing its nose towards the object and engaging in active exploration behaviors such as sniffing. Sitting, standing, or rearing next to the object without any sign of active exploration towards the object was not counted. The time a rat spent exploring each object during the retrieval test was converted into a discrimination ratio according to the general formula: (exploration time for novel object-location – exploration time for familiar object-location) / (exploration time for novel object-location + exploration time for familiar object-location). A value of zero indicates no exploration preference, whereas a positive value indicates preferential exploration of the displaced object, thus indicating memory of the familiar configuration. Additionally, the total time of object exploration (across both objects) and distance traveled were determined.

### Surgical implantation of cannulae and optic fibers

Before surgery, rats received an intraperitoneal injection of anesthetic mixture (0.005 mg/kg fentanyl, 2 mg/kg midazolam, and 0.15 mg/kg medetomidine). The surgery was carried out under general isoflurane anaesthesia (induction: 4%, maintenance: 0.8–1.25% in 0.35 l/min O2). Rats were placed in the stereotaxic frame, and the skull was exposed. Depending on the experimental condition, stainless steel guide cannulae (Plastics One) were implanted either unilaterally above the lateral ventricle (AP −0.5 mm, ML +1.5 mm, and DV −2.3 mm, 22-gauge, 7 mm, Experiment 2), bilaterally above the dorsal hippocampal CA1 (AP −4.0 mm, ML ±2.8 mm, and DV −1.4 mm, with an 8° lateral tilt, 22/23-gauge, 7 mm, Experiment 3 and 4) or the RSD (AP −4.0 mm, ML ±0.75 mm, and DV 0 mm, 23-gauge double guide with 1.5 mm inter-cannula spacing, 5 mm, Experiment 6). All cannulae were secured to the skull using four bone screws and cold-polymerizing dental resin. Dummy cannulae (Plastics One) were kept in place and removed only during infusions (28-gauge, 7 mm; 28−30-gauge, 7 mm; 28-gauge, 5.5 mm with 0.5 mm protrusion from the guide, for Experiment 2, 3−4 and 6, respectively).

For the optogenetic inactivation experiment (Experiment 5), animals underwent two separate surgeries: one for virus injection and one for optic fiber implantation. Rats were anesthetized and prepared as described above for cannula implantation. During the first surgery, animals were bilaterally injected with 500 nL of viral vector (AAV5-hSyn-Jaws-KGC-GFP-ER2; Addgene Plasmid #65014; diluted 1:4 in sterile PBS; resulting titer: ∼1.75 x 1012 vg/mL) targeting the dorsal CA1 (AP: −4.0 mm, ML: ±2.4 mm, DV: −2.3 mm), to enable AAV-mediated expression of Jaws, a red-shifted light-activated chloride inward pump. Expression was driven by the human synapsin promoter, which supports robust transgene expression across neuronal populations but does not distinguish between excitatory and inhibitory neurons. The virus was delivered at 0.1 µL/min via a sharpened glass pipette (Wiretrol II, Drummond Scientific; tip diameter <25 µm), which remained in place for 10 min post-injection to prevent backflow, then was automatically withdrawn at 0.2 µm/s using a motorized micromanipulator (MP-285, Sutter Instruments). Craniotomies were sealed with silicone elastomer (Kwik-Cast, World Precision Instruments) and cold-polymerizing dental resin (Palapress, Kulzer), and the wound was sutured. The second surgery was performed 3– 4 weeks later, allowing sufficient time for viral expression. In this procedure, two optic fibers (400 µm diameter; Thorlabs) were implanted bilaterally above the injection sites at AP −4.0, ML ±2.4, and DV −1.8. The optic fibers were secured to the skull using four bone screws and cold-polymerizing dental resin. Animals were given at least 7 days to recover.

### Substance infusion

In the NPY-Y1R antagonist experiment (Experiment 2), rats received an intracerebro-ventricular (ICV) infusion of the NPY-Y1 receptor antagonist BIBO 3304 (Tocris Bioscience, 15 μg dissolved in 2 μL of 0.9% saline solution) or 0.9% saline solution (2 µL at 0.8 µL/min) immediately after the OPR encoding phase. For the Hippocampal and RSD inactivation experiments (Experiment 3, 4 and 6), rats received an intracerebral infusion of the GABA-A receptor agonist muscimol (Sigma; 0.5 µg dissolved in 0.5 µL of 0.9% saline per hemisphere) or 0.9% saline solution (0.5 and 0.3 µL, at 0.25 and 0.1 µL/min, for hippocampus and RSD respectively) following standardized procedures (Sawangjit et al., 2022), either immediately after encoding (Experiment 3) or 15 min prior to the retrieval test (Experiment 4 and 6). Infusion was delivered using an automated syringe pump (PHD ULTRA, Harvard Apparatus). For substance administration, the injection cannula was connected to a 5-µL Hamilton microsyringe (Hamilton Company, Reno, NV) via polyethylene tubing. The injection cannula (28-, 30-gauge) protruded 2, 1 and 0.5 mm beyond the guide cannula for Experiment 2, 3−4 and 6, respectively. Following infusion, the cannulae were left in place for an additional minute to minimize backflow.

### Optogenetic inhibition of dorsal hippocampus

For optogenetic inhibition experiment (Experiment 5), during the encoding phase, light was delivered through fiber patch cables connected to a red-light LED source (625 nm, Fiber-Coupled, Thorlabs). The light output was regulated by a dedicated LED driver (T-Cube^TM^, Thorlabs) and controlled by an Arduino-based timing system programmed to generate the stimulation pattern. Light power at the fiber tip was calibrated before each session using a photodiode power meter and adjusted to 15mW to ensure effective neuronal inhibition while avoiding tissue heating. Continuous illumination was applied during the designated inhibition period (encoding phase) using a repeated cycle of 8 s ON followed by 2–2.5 s OFF, repeated for 10 min. The OFF period was slightly jittered to avoid rhythmic entrainment and neural artifacts. Patch cables were connected to the implanted optic-fiber ferrules immediately prior to the start of the trial and the system was switched on before placing the animal into the arena, ensuring consistent illumination from the beginning of the behavioral session.

### Histology

Correct placement of the cannulae, verification of viral expression, and - in the RSD inactivation experiment - localization of muscimol spread were assessed histologically after completion of the experiments. In the retrosplenial cohort, fluorescent muscimol (Hello Bio, 0.3 μg dissolved in 0.3 μL of 0.9% saline solution) was infused 20 minutes prior to perfusion to visualize the diffusion of the drug. All rats were perfused intracardially with 0.9% phosphate buffered saline (PBS) followed by 4% paraformaldehyde (PFA). After decapitation, brains were removed and post-fixed in 4% PFA for at least 24 h. Brains were stored in PBS at 4 °C until further processing. Coronal sections (70 µm) were cut on a vibratome (Microm HM 650 V, Thermo Fisher Scientific MICROM GmbH, Germany). In animals injected with the virus and in those infused with fluorescent muscimol, the intrinsic fluorescence signal was used to assess the extent and localization of viral expression and muscimol spread using an epifluorescence microscope (DMi8, Leica Microsystems, Germany).

### c-Fos immunocytochemistry

c-Fos immunocytochemistry was done as previously described^69^. For the comparison between experimental groups, c-Fos levels were normalized to those of a home cage control group which stayed in their home cage during the OPR task phases but, otherwise received the same treatment as the Wake-Hunger animals. In the main experiment, animals were transcardially perfused 90 minutes after the end of the retrieval phase (or corresponding interval of the Home Cage control group). The 90-min interval coincides with the peak of c-Fos protein expression following neuronal activation^70^. All further analyses were done by a blinded experimenter who was unaware of the individual rat’s experimental condition. Coronal slices (70 µm) at the respective bregma levels (see below) were cut with a vibratome. For immunostaining, free-floating sections were blocked with PBS containing 10 % horse serum and 0.3 % Triton-X for 90 min. Afterwards, the sections were incubated with 0.2 % Triton-X in PBS containing rabbit polyclonal anti-c-Fos primary antibody (1:1000 dilutions; Synaptic Systems, Germany, #226 003) for 48 h at 4 °C. Sections were washed with PBS (four times), then incubated with 0.2 % Triton-X in PBS containing donkey anti-rabbit Alexa Fluor 555 secondary antibody (1:1000 dilution; ThermoFisher Scientific, MA) for 24 h at 4 °C. Samples were incubated with NeuroTrace (1:200 dilutions; ThermoFisher Scientific), washed with PBS, mounted on gel-coated slides, and coverslipped with Vectashield antifade mounting medium (Vector Laboratories, CA).

Regions of interest (ROIs) were defined based on the literature about cortical, hippocampal, thalamic, and hypothalamic regions known to be involved in the formation of spatial memories and anatomically determined according to the rat brain atlas^71^. ROIs and their corresponding distances from bregma were: +3.24 mm for the prelimbic (PL), infralimbic (IL), and anterior cingulate (CG) cortices; −3.96 mm for the retrosplenial degranular (RSD) and granular (RSG) cortices, the parietal (PAR), perirhinal (PRh) and entorhinal (Ent) cortices, as well as the hippocampal cornu ammonis fields (CA1 and CA3) and the dentate gyrus (DG); −1.80 mm for the thalamic nucleus reuniens (RE) and reticular thalamic nucleus (RTN); and −2.50 mm for the lateral hypothalamus (LH) and the arcuate nucleus (Arc).

Images were obtained using a confocal laser scanning microscope (LSM 710; Carl Zeiss, Germany) with a low magnification (plan-apochromat oil-immersion objective, 25 ×/0.8NA; Z stack, 11 slices (50 µm thickness); scaling (per pixel), 0.33 µm × 0.33 µm × 3.5 µm). Settings for laser intensity, gain, offset and pinhole were optimized initially and held constant throughout the experiment. Automated c-Fos expressing cell counts were performed using CellProfiler software (Broad Institute, USA). Counting was done on z-stacks of the entire slice volume presented as maximum intensity projection. The sizes of the counting frames were 924,800 µm2 for LH, 693,600 µm2 for PL, 616,148 µm2 for CA1 and CA3, 578,200 µm2 for CG, 462,400 µm2 for IL, Arc and PRh, 346,800 µm2 for RE, RTN, DG, PAR and RSD, 231,200 µm2 for RSG and Ent.

### Data reduction and statistical analyses

A total of seven rats were excluded from data analyses for the following reasons: implant loss (n = 2; one from each Experiment 4 and 5), cannula misplacement (n = 4; two from Experiment 2, one from each Experiment 3 and 4), and implant occlusion (n = 1; from Experiment 4).

Statistical analyses were performed using R Statistical Software (R Core Team, 2021). Behavioral data from the OPR task were analyzed using analyses of variance (ANOVAs) that included either a Group factor (main experiment, NPY-Y1R antagonist experiment) or, in the remaining experiments, a Condition factor to reflect the Sleep-Fed and Wake-Hungry conditions, and a Minute factor reflecting the 5-min duration of the OPR retrieval phase. Analyses focused primarily on minute 1 and 3 of the retrieval phase which represent the most sensitive window for detecting OPR memory effects^18,19^, and were additionally performed across the full 5-min retrieval interval. To further specify significant main and interaction effects, global ANOVAs were followed by sub-ANOVAs performed pairwise between groups and, in case of significant Minute effects, separately for minute 1 and 3, and by two-sided post-hoc *t*-tests (Bonferroni correction). Discrimination indexes were also compared against chance level (µ = 0) using one-sample *t*-tests. c-Fos expression data were first normalized to the median value of the home-cage control group to account for inter-batch variability. Group differences were assessed using Aligned Rank Transform (ART) ANOVA. Additionally, c-Fos levels were compared with home-cage controls (µ = 1) using a one-sample Wilcoxon signed-rank test. Functional-network analysis was performed using the R package *igraph*^72^ to extract measures of degree and betweenness centrality^73^. Group differences between these measures were assessed using non-parametric permutation-based network comparison tests (NCT) allowing robust inference on network-level metrics.

## Acknowledgments

We thank I. Sauter for technical support, M. Harkotte for assistance with the optogenetic setup, J. Fechner for providing the script for the NCT, and M. Hallschmid for proof-reading. This work was supported by grants to J.B. from the Deutsche Forschungsgemeinschaft (FOR 5434) and the European Research Council (ERC AdG 883098 Sleep Balance). M.I. was supported by the Hertie Foundation (Hertie Network of Excellence in Clinical Neuroscience).

## Author contributions

M.I., J.B. and A.S. conceived the project. E.T., J.B. and A.S. designed the experiments. J.B. and A.S. supervised the project. Jo.B. A.O.G. and J.L.V. conducted extensive pilot experiments. E.T. and A.S. conducted the main experiments. S.D. conducted immunocytochemistry analyses. E.T. analyzed the data and drafted the manuscript. M.I., J.B., and A.S. edited and reviewed the manuscript. All authors approved the final version of the paper.

---

**Extended Data Fig. 1.**
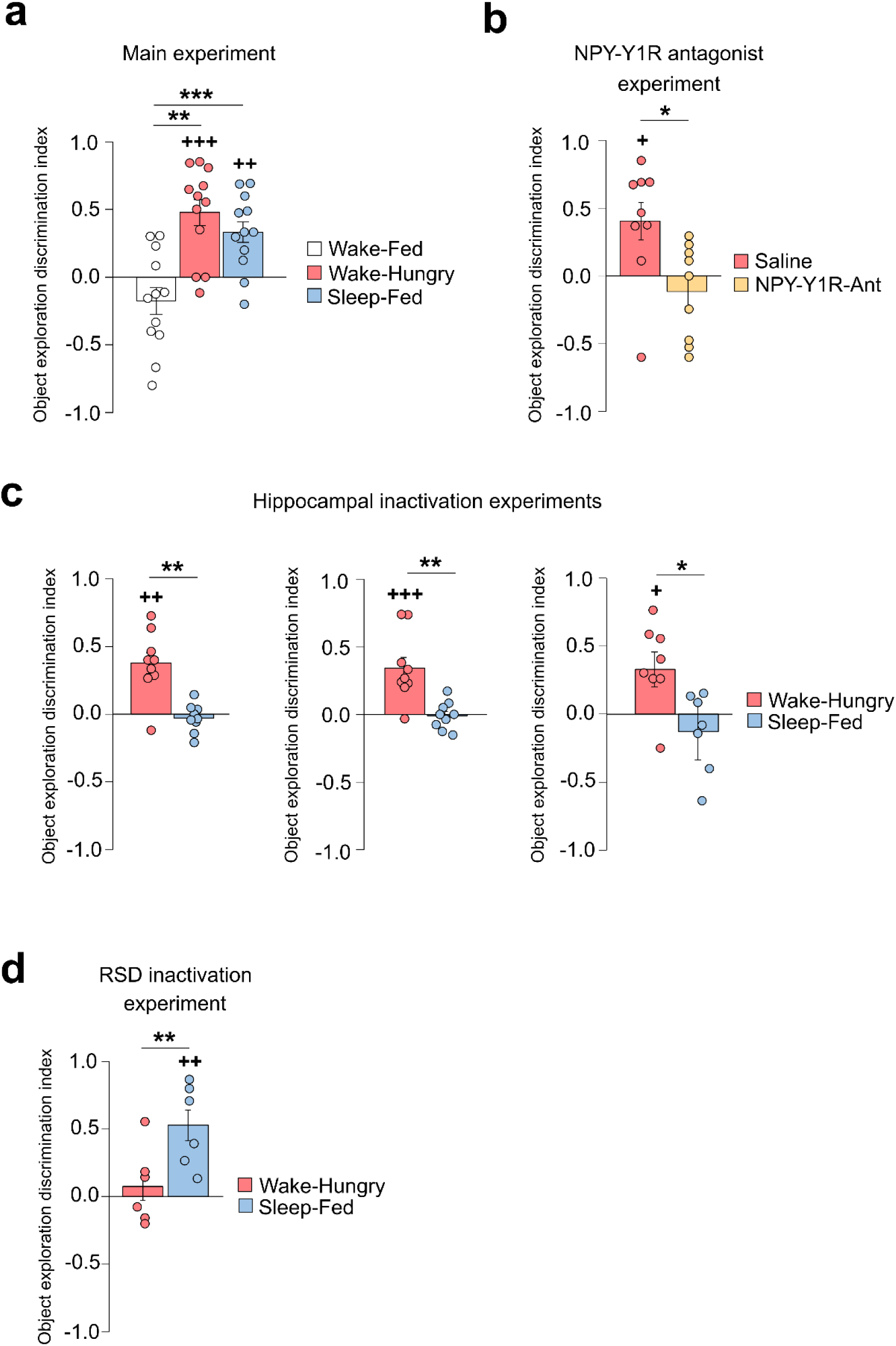
OPR discrimination indexes (DIs) across the entire 5-minute retrieval phase in the different experiments. **a**, Main experiment: Wake-Hungry (red) and Sleep-Fed (blue) groups showed significant OPR memory, whereas OPR memory was absent in the Wake-Fed group (white, F(2,33) = 12.82, *p* < 0.001, for Group main effect; n = 12 rats per group). **b**, NPY-Y1R antagonist experiment: OPR memory was absent in the Wake-Hungry rats of the NPY-Y1R-Ant group (light gold, n = 9 rats) but was preserved in the Saline group (red, n = 9 rats). **c**, Hippocampal inactivation experiments: reversible CA1 inactivation during consolidation (*left,* n = 9 rats), retrieval (*center*, n = 9 rats) and encoding (*right*, n = 7 rats) phases abolished OPR memory in Sleep-Fed conditions (blue) whereas OPR memory was preserved in Wake-Hungry conditions (red). **d**, RSD inactivation experiment (n = 6 rats): Reversible inactivation of RSD during OPR retrieval phase abolished OPR memory in Wake-Hungry (red) but not Sleep-Fed (blue) conditions. Data are shown as mean ± s.e.m. with overlaid dot plots. \*\*\**p* < 0.001, \*\**p* < 0.05, \**p* < 0.05 for pairwise two-sided *t*-test. ^+++^*p* < 0.001, ^++^*p* < 0.01, ^+^*p* < 0.05, for one-sample *t*-tests against chance level.

**Extended Data Fig. 2.**
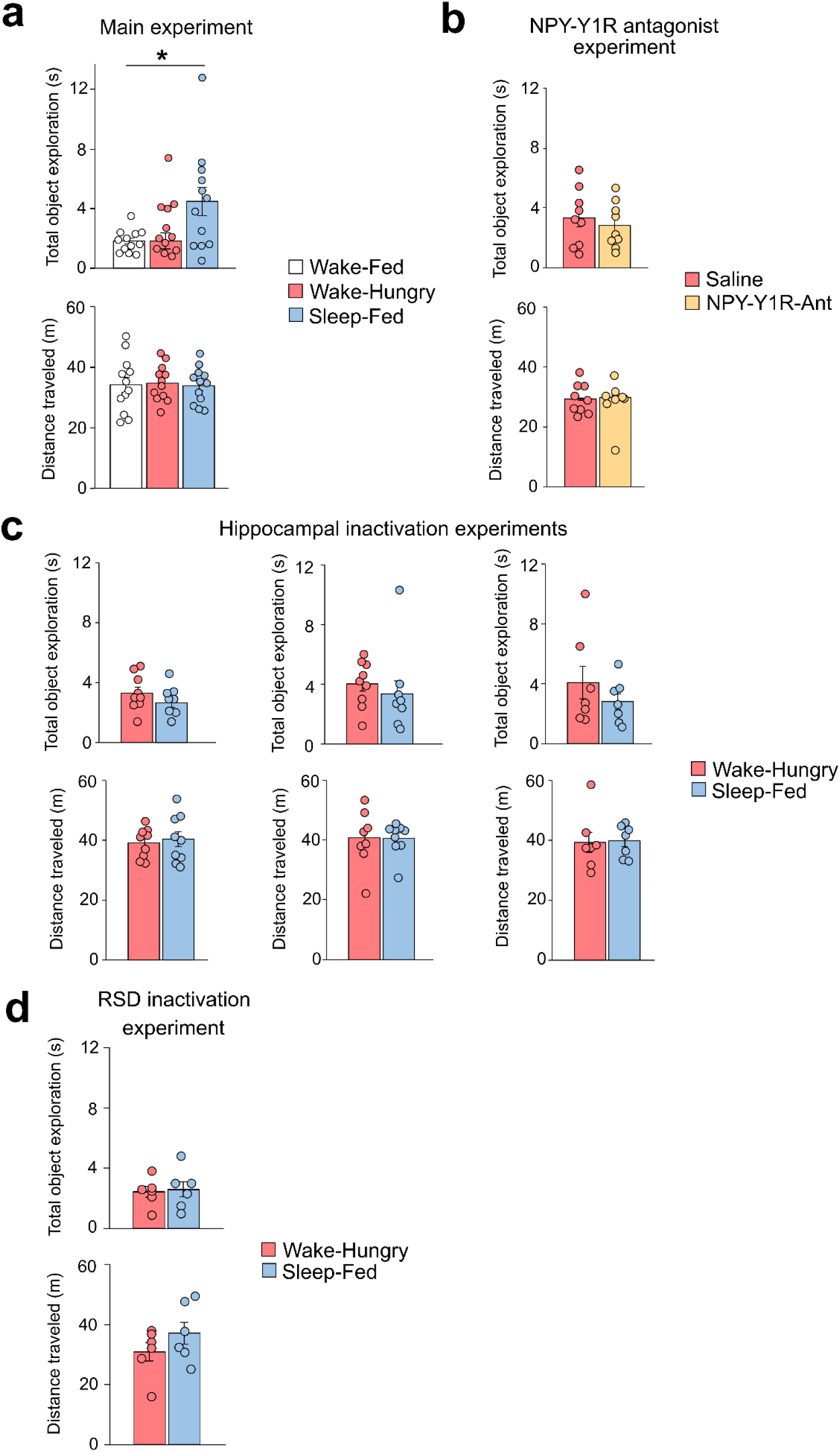
Total object exploration time and distance traveled across the entire 5-minute retrieval phase in the different experiments. Control measures of total object exploration time (*top*) and distance traveled (*bottom*) during the OPR retrieval phase in (a) the main experiment, (b) NPY-Y1R antagonist experiment, (c) hippocampal inactivation experiments, and (d) RSD inactivation experiment. There were no significant differences between experimental conditions in any of the experiments, except in the main experiment. Data are showed as mean ± s.e.m. with overlaid dot plots*. *p < 0.05, for pairwise two-sided t-tests*.

**Extended Data Fig. 3.**
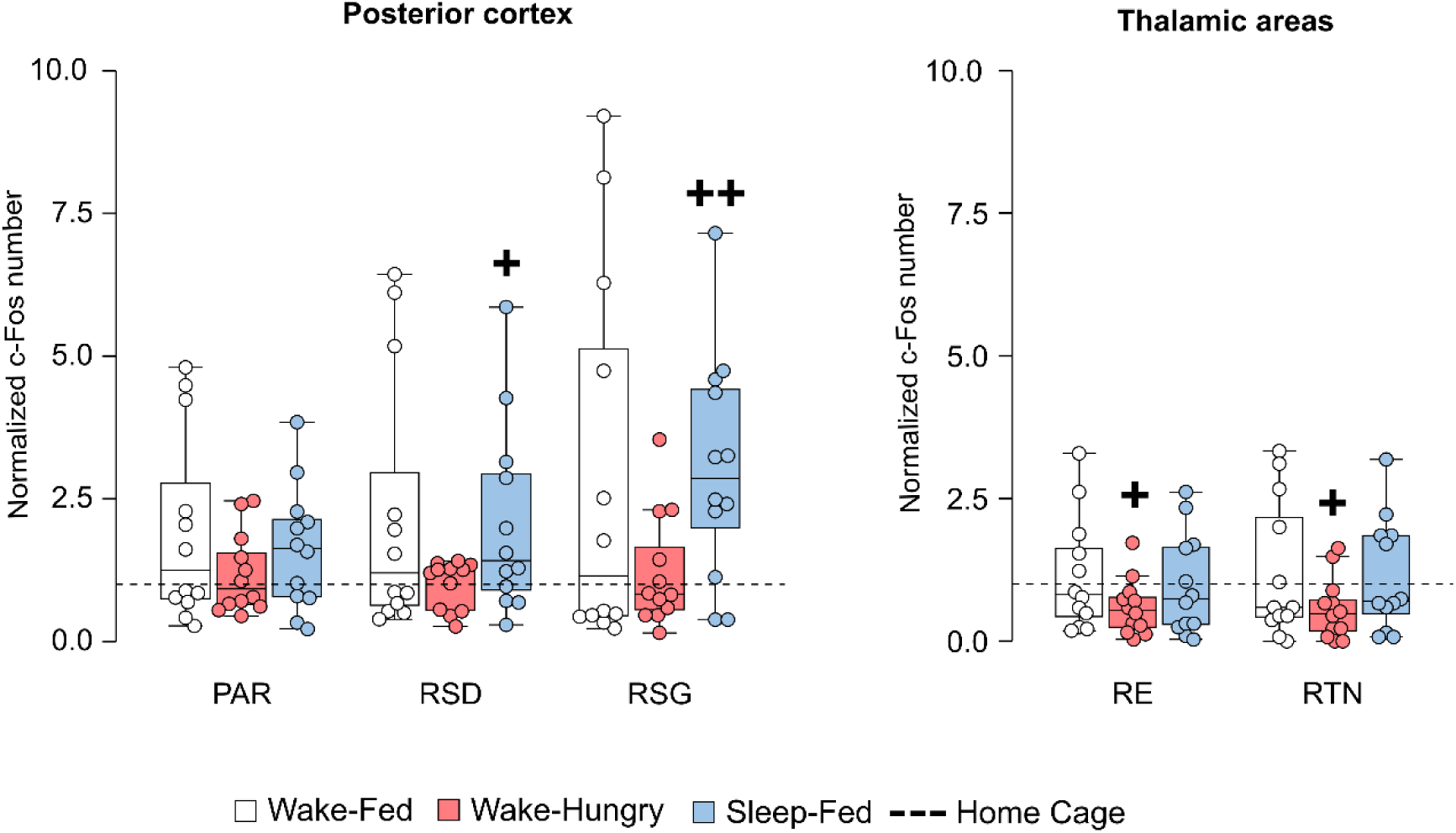
c-Fos expression during OPR memory retrieval – posterior cortex and thalamic areas. Number of c-Fos positive cells relative to home cage control group (set to 1, dashed lines) at OPR retrieval phase in *(left)* posterior cortex and (*right*) thalamic areas in the Wake-Fed (white), Wake-Hungry (red), and Sleep-Fed (blue) groups of the main experiment (n = 12 rats per group). Data shown as boxplots with overlaid dot plots. Boxes represent the median and interquartile range (IQR), whiskers extend to 1.5 x IQR. There were no significant differences between groups. ^++^*p* < 0.01, ^+^*p* < 0.05, for a one-sample Wilcoxon signed-rank test against home cage control. Abbreviations: PAR = parietal cortex, RE = nucleus reuniens, RSD = dysgranular retrosplenial cortex, RSG = granular retrosplenial cortex, RTN = Reticular nucleus of the thalamus.

**Extended Data Fig. 4.**
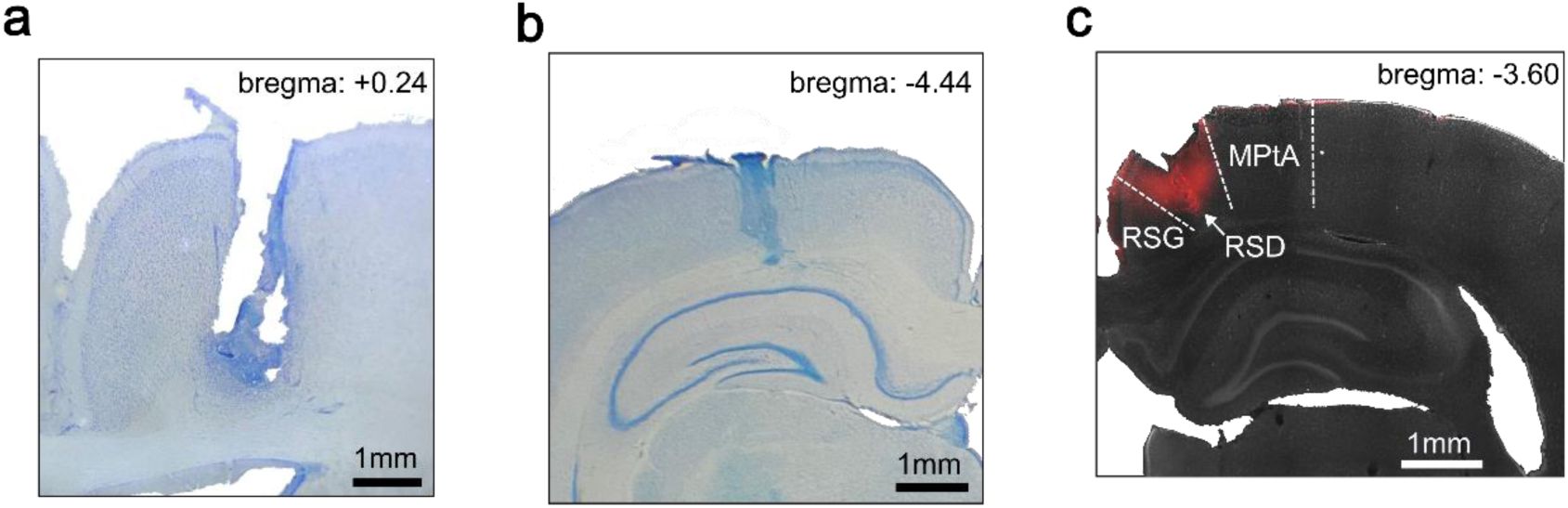
Histological verification of cannula placement and muscimol diffusion. **a**, Representative histological image showing the placement of the cannula used for intracerebroventricular infusion in the NPY-Y1R antagonist experiment. The guide cannula was positioned above the lateral ventricle, with the internal cannula extending 2 mm beyond the guide to reach the lateral ventricle. **b**, Cannula placement in the Hippocampal inactivation during consolidation and retrieval experiments. The guide cannula was positioned above the dorsal hippocampus, with the internal cannula extending 1 mm to target the CA1. **c**, Muscimol diffusion in the dysgranular retrosplenial cortex (RSD) inactivation experiment. Fluorescence-labeled muscimol (red) was infused prior to perfusion to visualize drug spread. Muscimol diffused into the RSD with minimal spread into the adjacent granular retrosplenial cortex (RSG). MPtA = medial parietal association cortex.

## Notes

### Competing Interest Statement

The authors have declared no competing interest.

## References

1. Müller, G.E., and Pilzecker, A. (1900). Experimentelle Beiträge zur Lehre vom Gedächtniss / von G.E. Müller und A. Pilzecker. Wellcome Collection. https://wellcomecollection.org/works/d68pp8k8.

2. Moscovitch, M., Rosenbaum, R.S., Gilboa, A., Addis, D.R., Westmacott, R., Grady, C., McAndrews, M.P., Levine, B., Black, S., Winocur, G., et al. (2005). Functional neuroanatomy of remote episodic, semantic and spatial memory: a unified account based on multiple trace theory. Journal of Anatomy 207, 35–66. 10.1111/j.1469-7580.2005.00421.x.

3. Tarder-Stoll, H., Sekeres, M.J., Levine, B., and Moscovitch, M. (2025). Adaptive episodic memory: How multiple memory representations drive behaviour in humans and non-humans. Physiological Reviews. 10.1152/physrev.00005.2025.

4. Alexander, A.S., Place, R., Starrett, M.J., Chrastil, E.R., and Nitz, D.A. (2023). Rethinking retrosplenial cortex: Perspectives and predictions. Neuron 111, 150–175. 10.1016/j.neuron.2022.11.006.

5. Diekelmann, S., and Born, J. (2010). The memory function of sleep. Nat Rev Neurosci 11, 114–126. 10.1038/nrn2762.

6. Lutz, N.D., Harkotte, M., and Born, J. (2026). Sleep’s contribution to memory formation. Physiol Rev 106, 363–483. 10.1152/physrev.00054.2024.

7. Heine, R. (1914). Über Wiedererkennen und rückwirkende Hemmung (Johann Ambrosius Barth).

8. Sawangjit, A., Oyanedel, C.N., Niethard, N., Salazar, C., Born, J., and Inostroza, M. (2018). The hippocampus is crucial for forming non-hippocampal long-term memory during sleep. Nature 564, 109–113. 10.1038/s41586-018-0716-8.

9. Brodt, S., Inostroza, M., Niethard, N., and Born, J. (2023). Sleep—A brain-state serving systems memory consolidation. Neuron 111, 1050–1075. 10.1016/j.neuron.2023.03.005.

10. Gridchyn, I., Schoenenberger, P., O’Neill, J., and Csicsvari, J. (2020). Assembly-Specific Disruption of Hippocampal Replay Leads to Selective Memory Deficit. Neuron 106, 291–300.e6. 10.1016/j.neuron.2020.01.021.

11. Schapiro, A.C., Reid, A.G., Morgan, A., Manoach, D.S., Verfaellie, M., and Stickgold, R. (2019). The hippocampus is necessary for the consolidation of a task that does not require the hippocampus for initial learning. Hippocampus 29, 1091–1100. 10.1002/hipo.23101.

12. Sawangjit, A., Harkotte, M., Oyanedel, C.N., Niethard, N., Born, J., and Inostroza, M. (2022). Two distinct ways to form long-term object recognition memory during sleep and wakefulness. Proceedings of the National Academy of Sciences 119, e2203165119. 10.1073/pnas.2203165119.

13. Chouhan, N.S., Griffith, L.C., Haynes, P., and Sehgal, A. (2021). Availability of food determines the need for sleep in memory consolidation. Nature 589, 582–585. 10.1038/s41586-020-2997-y.

14. Fadda, M., Hasakiogullari, I., Temmerman, L., Beets, I., Zels, S., and Schoofs, L. (2019). Regulation of Feeding and Metabolism by Neuropeptide F and Short Neuropeptide F in Invertebrates. Front. Endocrinol. 10. 10.3389/fendo.2019.00064.

15. Barker, G.R.I., and Warburton, E.C. (2011). When Is the Hippocampus Involved in Recognition Memory? J. Neurosci. 31, 10721–10731. 10.1523/JNEUROSCI.6413-10.2011.

16. Chao, O.Y., de Souza Silva, M.A., Yang, Y.-M., and Huston, J.P. (2020). The medial prefrontal cortex - hippocampus circuit that integrates information of object, place and time to construct episodic memory in rodents: behavioral, anatomical and neurochemical properties. Neurosci Biobehav Rev 113, 373–407. 10.1016/j.neubiorev.2020.04.007.

17. Marks, J.L., Li, M., Schwartz, M., Porte, D., and Baskin, D.G. (1992). Effect of fasting on regional levels of neuropeptide Y mRNA and insulin receptors in the rat hypothalamus: An autoradiographic study. Molecular and Cellular Neuroscience 3, 199–205. 10.1016/1044-7431(92)90039-5.

18. Dix, S.L., and Aggleton, J.P. (1999). Extending the spontaneous preference test of recognition: evidence of object-location and object-context recognition. Behavioural Brain Research 99, 191–200. 10.1016/S0166-4328(98)00079-5.

19. Ozawa, T., Yamada, K., and Ichitani, Y. (2011). Long-term object location memory in rats: effects of sample phase and delay length in spontaneous place recognition test. Neurosci Lett 497, 37–41. 10.1016/j.neulet.2011.04.022.

20. Sawangjit, A., Oyanedel, C.N., Niethard, N., Born, J., and Inostroza, M. (2020). Deepened sleep makes hippocampal spatial memory more persistent. Neurobiology of Learning and Memory 173, 107245. 10.1016/j.nlm.2020.107245.

21. Inostroza, M., Binder, S., and Born, J. (2013). Sleep-dependency of episodic-like memory consolidation in rats. Behav Brain Res 237, 15–22. 10.1016/j.bbr.2012.09.011.

22. Ishikawa, H., Yamada, K., Pavlides, C., and Ichitani, Y. (2014). Sleep deprivation impairs spontaneous object-place but not novel-object recognition in rats. Neurosci Lett 580, 114–118. 10.1016/j.neulet.2014.08.004.

23. Oyanedel, C.N., Binder, S., Kelemen, E., Petersen, K., Born, J., and Inostroza, M. (2014). Role of slow oscillatory activity and slow wave sleep in consolidation of episodic-like memory in rats. Behav Brain Res 275, 126–130. 10.1016/j.bbr.2014.09.008.

24. Wieland, H.A., Engel, W., Eberlein, W., Rudolf, K., and Doods, H.N. (1998). Subtype selectivity of the novel nonpeptide neuropeptide Y Y1 receptor antagonist BIBO 3304 and its effect on feeding in rodents. Br J Pharmacol 125, 549–555. 10.1038/sj.bjp.0702084.

25. Chambers, A.P., and Woods, S.C. (2012). The Role of Neuropeptide Y in Energy Homeostasis. In Appetite Control, H.-G. Joost, ed. (Springer), pp. 23–45. 10.1007/978-3-642-24716-3_2.

26. Finsterwald, C., and Alberini, C.M. (2014). Stress and glucocorticoid receptor-dependent mechanisms in long-term memory: from adaptive responses to psychopathologies. Neurobiol Learn Mem 112, 17–29. 10.1016/j.nlm.2013.09.017.

27. Roy, D.S., Park, Y.-G., Kim, M.E., Zhang, Y., Ogawa, S.K., DiNapoli, N., Gu, X., Cho, J.H., Choi, H., Kamentsky, L., et al. (2022). Brain-wide mapping reveals that engrams for a single memory are distributed across multiple brain regions. Nat Commun 13, 1799. 10.1038/s41467-022-29384-4.

28. Chao, O.Y., Nikolaus, S., Yang, Y.-M., and Huston, J.P. (2022). Neuronal circuitry for recognition memory of object and place in rodent models. Neurosci Biobehav Rev 141, 104855. 10.1016/j.neubiorev.2022.104855.

29. Berrios, J., Li, C., Madara, J.C., Garfield, A.S., Steger, J.S., Krashes, M.J., and Lowell, B.B. (2021). Food cue regulation of AGRP hunger neurons guides learning. Nature 595, 695–700. 10.1038/s41586-021-03729-3.

30. Tanimizu, T., Kenney, J.W., Okano, E., Kadoma, K., Frankland, P.W., and Kida, S. (2017). Functional Connectivity of Multiple Brain Regions Required for the Consolidation of Social Recognition Memory. J Neurosci 37, 4103–4116. 10.1523/JNEUROSCI.3451-16.2017.

31. Roux, L., Hu, B., Eichler, R., Stark, E., and Buzsáki, G. (2017). Sharp wave ripples during learning stabilize the hippocampal spatial map. Nat Neurosci 20, 845–853. 10.1038/nn.4543.

32. López, A.J., Kramár, E., Matheos, D.P., White, A.O., Kwapis, J., Vogel-Ciernia, A., Sakata, K., Espinoza, M., and Wood, M.A. (2016). Promoter-Specific Effects of DREADD Modulation on Hippocampal Synaptic Plasticity and Memory Formation. J Neurosci 36, 3588–3599. 10.1523/JNEUROSCI.3682-15.2016.

33. Eichenbaum, H. (2000). A cortical-hippocampal system for declarative memory. Nat Rev Neurosci 1, 41–50. 10.1038/35036213.

34. Moser, M.-B., and Moser, E.I. (1998). Distributed Encoding and Retrieval of Spatial Memory in the Hippocampus. J. Neurosci. 18, 7535–7542. 10.1523/JNEUROSCI.18-18-07535.1998.

35. Teuber, H.L. (1955). Physiological psychology. Annu Rev Psychol 6, 267–296. 10.1146/annurev.ps.06.020155.001411.

36. Bisby, J., and Burgess, N. (2017). Differential effects of negative emotion on memory for items and associations, and their relationship to intrusive imagery. Current Opinion in Behavioral Sciences 17, 124–132. 10.1016/j.cobeha.2017.07.012.

37. Poldrack, R.A., Clark, J., Paré-Blagoev, E.J., Shohamy, D., Creso Moyano, J., Myers, C., and Gluck, M.A. (2001). Interactive memory systems in the human brain. Nature 414, 546–550. 10.1038/35107080.

38. Schwabe, L., and Wolf, O.T. (2013). Stress and multiple memory systems: from “thinking” to “doing.” Trends Cogn Sci 17, 60–68. 10.1016/j.tics.2012.12.001.

39. Squire, L.R., and Zola, S.M. (1996). Structure and function of declarative and nondeclarative memory systems. Proc Natl Acad Sci U S A 93, 13515–13522. 10.1073/pnas.93.24.13515.

40. Nguyen, N.D., Tucker, M.A., Stickgold, R., and Wamsley, E.J. (2013). Overnight Sleep Enhances Hippocampus-Dependent Aspects of Spatial Memory. Sleep 36, 1051–1057. 10.5665/sleep.2808.

41. Simon, K.C., Clemenson, G.D., Zhang, J., Sattari, N., Shuster, A.E., Clayton, B., Alzueta, E., Dulai, T., de Zambotti, M., Stark, C., et al. (2022). Sleep facilitates spatial memory but not navigation using the Minecraft Memory and Navigation task. Proc Natl Acad Sci U S A 119, e2202394119. 10.1073/pnas.2202394119.

42. Bacci, A., Huguenard, J.R., and Prince, D.A. (2002). Differential modulation of synaptic transmission by neuropeptide Y in rat neocortical neurons. Proc Natl Acad Sci U S A 99, 17125–17130. 10.1073/pnas.012481899.

43. Kornhuber, J., and Zoicas, I. (2017). Neuropeptide Y prolongs non-social memory and differentially affects acquisition, consolidation, and retrieval of non-social and social memory in male mice. Sci Rep 7, 6821. 10.1038/s41598-017-07273-x.

44. Totani, Y., Nakai, J., Dyakonova, V.E., Lukowiak, K., Sakakibara, M., and Ito, E. (2020). Induction of LTM following an Insulin Injection. eNeuro 7, ENEURO.0088-20.2020. 10.1523/ENEURO.0088-20.2020.

45. Verma, D., Wood, J., Lach, G., Herzog, H., Sperk, G., and Tasan, R. (2016). Hunger Promotes Fear Extinction by Activation of an Amygdala Microcircuit. Neuropsychopharmacology 41, 431–439. 10.1038/npp.2015.163.

46. Yang, X., Miao, X., Schweiggart, F., Großmann, S., Rauss, K., Hallschmid, M., Born, J., and Lutz, N.D. (2025). The effect of fasting on human memory consolidation. Neurobiology of Learning and Memory 218, 108034. 10.1016/j.nlm.2025.108034.

47. Beck, B. (2006). Neuropeptide Y in normal eating and in genetic and dietary-induced obesity. Philos Trans R Soc Lond B Biol Sci 361, 1159–1185. 10.1098/rstb.2006.1855.

48. Roozendaal, B. (2002). Stress and Memory: Opposing Effects of Glucocorticoids on Memory Consolidation and Memory Retrieval. Neurobiology of Learning and Memory 78, 578–595. 10.1006/nlme.2002.4080.

49. Schwabe, L., Hermans, E.J., Joëls, M., and Roozendaal, B. (2022). Mechanisms of memory under stress. Neuron 110, 1450–1467. 10.1016/j.neuron.2022.02.020.

50. Melo, I., and Ehrlich, I. (2016). Sleep supports cued fear extinction memory consolidation independent of circadian phase. Neurobiol Learn Mem 132, 9–17. 10.1016/j.nlm.2016.04.007.

51. Dietze, S., Lees, K.R., Fink, H., Brosda, J., and Voigt, J.-P. (2016). Food Deprivation, Body Weight Loss and Anxiety-Related Behavior in Rats. Animals (Basel) 6, 4. 10.3390/ani6010004.

52. Heilig, M. (2004). The NPY system in stress, anxiety and depression. Neuropeptides 38, 213–224. 10.1016/j.npep.2004.05.002.

53. Reichmann, F., and Holzer, P. (2016). Neuropeptide Y: A stressful review. Neuropeptides 55, 99–109. 10.1016/j.npep.2015.09.008.

54. Girardeau, G., Benchenane, K., Wiener, S.I., Buzsáki, G., and Zugaro, M.B. (2009). Selective suppression of hippocampal ripples impairs spatial memory. Nat Neurosci 12, 1222–1223. 10.1038/nn.2384.

55. Danieli, K., Guyon, A., and Bethus, I. (2023). Episodic Memory formation: A review of complex Hippocampus input pathways. Progress in Neuro-Psychopharmacology and Biological Psychiatry 126, 110757. 10.1016/j.pnpbp.2023.110757.

56. Preston, A.R., and Eichenbaum, H. (2013). Interplay of Hippocampus and Prefrontal Cortex in Memory. Current Biology 23, R764–R773. 10.1016/j.cub.2013.05.041.

57. O’Keefe, J. (1978). The Hippocampus as a Cognitive Map (Oxford university press).

58. Vorhees, C.V., and Williams, M.T. (2014). Assessing Spatial Learning and Memory in Rodents. ILAR J 55, 310–332. 10.1093/ilar/ilu013.

59. Oliveira, A.M.M., Hawk, J.D., Abel, T., and Havekes, R. (2010). Post-training reversible inactivation of the hippocampus enhances novel object recognition memory. Learn Mem 17, 155–160. 10.1101/lm.1625310.

60. Li, Y., Ren, M., Liu, B., Jiang, T., Jia, X., Zhang, H., Gong, H., and Wang, X. (2025). Dissection of the long-range circuit of the mouse intermediate retrosplenial cortex. Commun Biol 8, 56. 10.1038/s42003-025-07463-8.

61. Miyashita, T., and Rockland, K.S. (2007). GABAergic projections from the hippocampus to the retrosplenial cortex in the rat. European Journal of Neuroscience 26, 1193–1204. 10.1111/j.1460-9568.2007.05745.x.

62. Balcerek, E., Włodkowska, U., and Czajkowski, R. (2024). FOS mapping reveals two complementary circuits for spatial navigation in mouse. Sci Rep 14, 21252. 10.1038/s41598-024-72272-8.

63. Hindley, E.L., Nelson, A.J.D., Aggleton, J.P., and Vann, S.D. (2014). The rat retrosplenial cortex is required when visual cues are used flexibly to determine location. Behavioural Brain Research 263, 98–107. 10.1016/j.bbr.2014.01.028.

64. Vann, S.D., and Aggleton, J.P. (2005). Selective dysgranular retrosplenial cortex lesions in rats disrupt allocentric performance of the radial-arm maze task. Behav Neurosci 119, 1682–1686. 10.1037/0735-7044.119.6.1682.

65. Schwarting, R.K.W., and Busse, S. (2017). Behavioral facilitation after hippocampal lesion: A review. Behavioural Brain Research 317, 401–414. 10.1016/j.bbr.2016.09.058.

66. Kolb, B., and Whishaw, I.Q. (1998). BRAIN PLASTICITY AND BEHAVIOR. Annual Review of Psychology 49, 43–64. 10.1146/annurev.psych.49.1.43.

67. Elias, A., Padinjakara, N., and Lautenschlager, N.T. (2023). Effects of intermittent fasting on cognitive health and Alzheimer’s disease. Nutr Rev 81, 1225–1233. 10.1093/nutrit/nuad021.

68. Parhizkar, S., and Holtzman, D.M. (2025). The night’s watch: Exploring how sleep protects against neurodegeneration. Neuron 113, 817–837. 10.1016/j.neuron.2025.02.004.

69. Dimitrov, S., Shan, X., Born, J., and Inostroza, M. (2026). Impact of tissue storage time on immunodetection of c-Fos and GAD67 in the rat brain. J Neurosci Methods 425, 110602. 10.1016/j.jneumeth.2025.110602.

70. Zangenehpour, S., and Chaudhuri, A. (2003). Differential induction and decay curves of c-fos and zif268 revealed through dual activity maps (vol 109, pg 221, 2002). Brain research. Molecular brain research 109, 221–225. 10.1016/S0169-328X(03)00165-7.

71. Paxinos, G., and Watson, C. (2013). The Rat Brain in Stereotaxic Coordinates 7th ed. (Academic Press).

72. Csárdi, G., Nepusz, T., Traag, V., Horvát, S., Zanini, F., Noom, D., Müller, K., Antonov, M., Initiative, C.Z., Schoch, D., et al. (2026). igraph: Network Analysis and Visualization. Version 2.2.2.

73. Brandes, U. (2005). Network Analysis: Methodological Foundations (Springer Science & Business Media).

